# Neural dynamics track the approach to physical exhaustion

**DOI:** 10.64898/2026.11.12.766118

**Authors:** Chiara Avancini, Juan E. Arco, Clara Alameda, Tristán A. Bekinschtein, Daniel Sanabria, Luis F. Ciria

## Abstract

Physical exhaustion has long been described through its cardiorespiratory and metabolic cost, while the neural dynamics of its approach remain largely unexplored. Here we recorded high-density electroencephalography in 28 trained cyclists during a maximal incremental test to characterise how neural dynamics change towards exhaustion. As physiological strain rose, oscillatory power shifted to lower frequencies, neural activity became more regular and predictable, and functional coupling extended to longer timescales. Participant-specific models built on neural dynamics tracked proximity to exhaustion with accuracy comparable to cardiorespiratory measures. Over the final minutes, neural features remained informative while cardiorespiratory measures lost accuracy. Models trained across participants still tracked proximity to exhaustion, but less precisely. The approach to exhaustion thus follows a common neural path, tuned to each individual.

## INTRODUCTION

Evolution has endowed humans with exceptional endurance capabilities, providing key survival and health advantages (*1, 2*). However, as exercise becomes more intense or is prolonged, even individuals with superior innate and trained capacities inevitably reach a limit where physical demand can no longer be sustained, forcing the cessation of exertion. This phenomenon stems from widespread changes across all physiological systems, constantly receiving and sending signals to the brain (*3, 4*), which constitutes the central focus of this study.

Using incremental and time-to-exhaustion exercise tests, exercise physiologists have extensively characterised changes in cardiovascular, metabolic, and muscular activity from rest to exhaustion, describing, for instance, the well-known aerobic and anaerobic metabolic pathways (*5*). Far less attention has been paid to the brain, despite its widely acknowledged role in the regulation of endurance performance (*6*). The few studies that have investigated brain functioning during high-intensity exercise (e.g., *7–10*) have focused almost exclusively on electroencephalography (EEG) frequency dynamics, reporting changes in the power of the canonical EEG bands as effort rises. Neural dynamics, however, are far richer than their spectrum. How activity unfolds over time, how regular it is and how it is coordinated across the cortex carry information that band power does not capture (e.g., *11*). A fine-grained characterisation of how these dynamics reorganise as exhaustion approaches is still lacking.

For decades, research on human performance has sought not only to describe exhaustion but also to anticipate it. These efforts have relied mainly on models based on cardiorespiratory and metabolic markers, whose estimates for a given individual remain imprecise (*12*). The brain, which integrates signals from across the body and drives the motor output that sustains exercise, is a natural candidate to provide the information that these models lack. Whether neural dynamics carry such information, or merely mirror cardiorespiratory strain, remains largely unexplored. Maximal incremental effort tests offer a well-suited experimental model to address this question, as effort builds up gradually over minutes yet ends in the individual reaching physical exhaustion, characterised by a sudden drop in performance due to an inability to sustain the required effort intensity, and the eventual cessation of exercise. Such a gradual build-up towards an abrupt end is characteristic of transitions between physiological states (e.g., from wakefulness to sleep), which are often preceded by a progressive reorganisation of the dynamics of the system involved (e.g., *13, 14*). If the approach to exhaustion follows the same course, neural dynamics should carry information about how close each individual is to exhaustion, and could complement cardiorespiratory markers, which become less informative near the limit (*15*).

Here we combined high-density EEG with cardiorespiratory measures in 28 trained amateur cyclists during a maximal incremental cycling test until exhaustion (Fig. 1A). We show that neural dynamics reorganise systematically as the limit approaches and track each participant’s proximity to exhaustion. When neural and cardiorespiratory measures are combined, the models track this proximity better than either alone. These findings suggest that neural dynamics carry information about the approach to the physical limit that cardiorespiratory measures do not capture.

**Figure 1.**
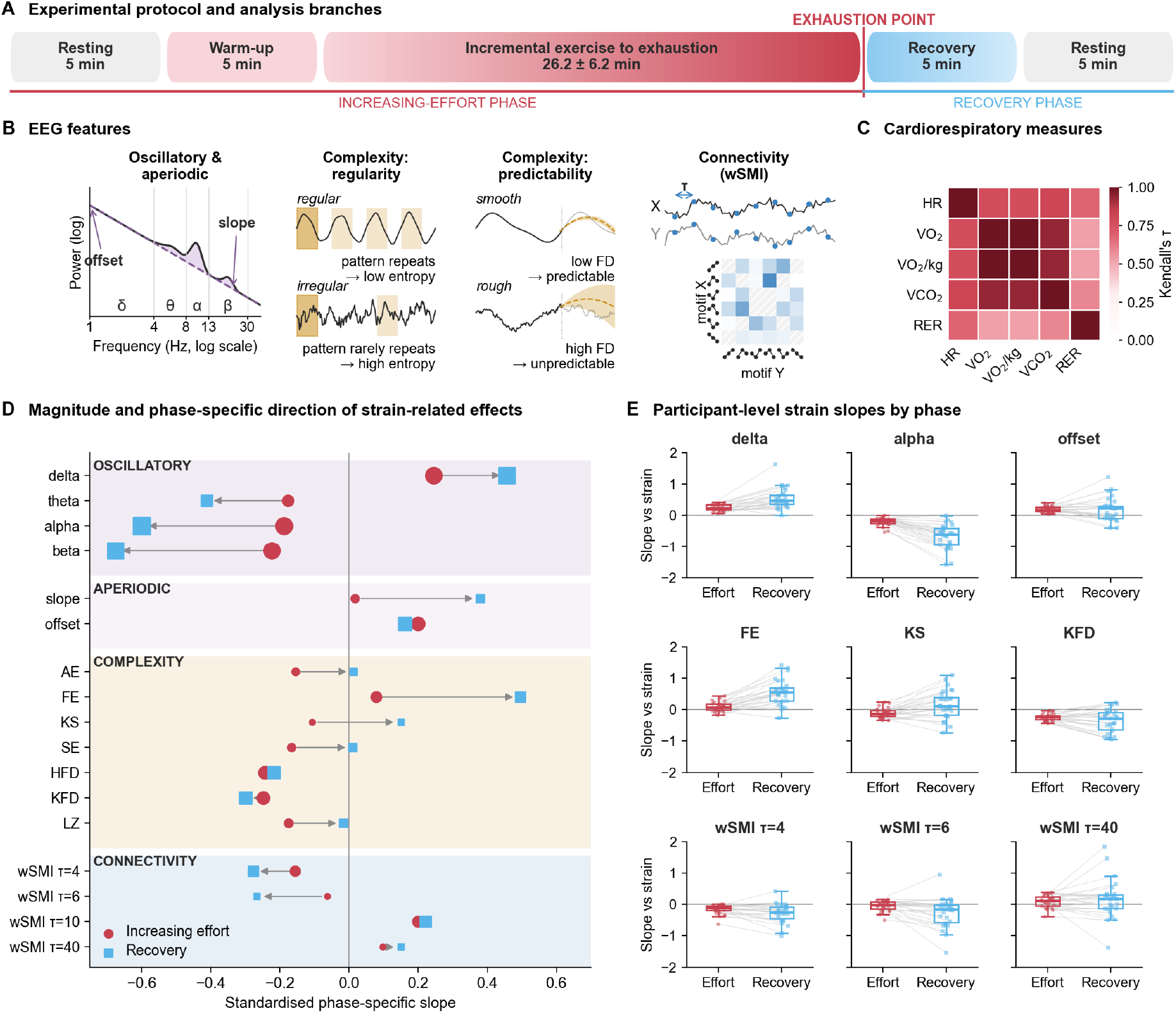
Physiological strain is coupled to a reversible reorganisation of neural activity. (A) Experimental protocol. After rest and a warm-up, participants (n = 28) cycled against an incremental load until exhaustion, then pedalled at low intensity and rested again. The protocol was split at exhaustion into an increasing-effort phase (red) and a recovery phase (blue). (B) The four families of EEG features, computed on consecutive 3-s epochs. The spectrum was decomposed with specparam into oscillatory power in the canonical bands and an aperiodic component described by its offset and slope. Following Lau et al. (*16*), complexity was indexed by regularity, which quantifies how often patterns recur (AE, SE, FE, LZ), and by predictability, which describes how the signal evolves over time (KS, HFD, KFD). Connectivity was estimated with weighted symbolic mutual information (wSMI) between ordinal motifs at four lags, discounting identical and opposite pairs (hatched) that are likely to arise from common sources. (C) Kendall’s τ between the cardiorespiratory measures, whose first principal component indexed physiological strain. (D) Strain-related effects for all the EEG features, from piecewise linear models with strain, phase and their interaction as predictors. Circles and squares depict the standardised slope on strain during increasing effort and recovery, and arrows run from the first to the second. Marker size is proportional to the effect size (ηp², Table S1). All effects survived family-wise correction. (E) Participant-level slopes on strain in each phase for nine representative features (two oscillatory, one aperiodic, three complexity and three wSMI features). Points are participants, lines join the two slopes of each participant, and boxes show the median and interquartile range. As strain increased, power shifted from theta, alpha and beta towards delta, the aperiodic offset rose, six of the seven complexity features decreased and wSMI coupling moved towards slower timescales. Most associations kept their sign as strain declined during recovery, even though muscular demand had dropped and time kept advancing.

## RESULTS

### Neural dynamics shift towards a slower regime as effort increases

To examine how neural dynamics changed across the full range of physiological demand, from rest to exhaustion and through recovery, we first extracted a set of EEG features that capture complementary aspects of these dynamics (i.e., oscillatory power, the aperiodic component of the power spectrum, signal complexity and functional connectivity; Fig. 1B). We then derived a single index of physiological demand from the cardiorespiratory measures recorded (oxygen uptake, body-mass-adjusted oxygen uptake, carbon dioxide output, respiratory exchange ratio and heart rate). As expected, all these measures were highly intercorrelated (Kendall’s τ = 0.526–1.000, Fig. 1C), and a principal component analysis yielded a first component that accounted for 87.5% of their variance, with positive loadings on all five measures (range = 0.33–0.49). We used this component, hereafter termed physiological strain, as the predictor in piecewise linear models fitted to each EEG feature, which estimated its association with strain separately for the increasing-effort and recovery phases.

We first fitted piecewise linear models relating physiological strain to oscillatory power in the canonical delta, theta, alpha and beta bands and to the offset and slope of the aperiodic 1/f component of the spectrum (*17*). Oscillatory power shifted towards slower frequencies as strain rose, with a significant effect in all four bands after family-wise correction (all p < 0.001, η_p_² = 0.240-0.548, Fig. 1D and Table S1). Delta power increased with physiological strain, whereas theta, alpha and beta power decreased. The aperiodic offset rose with strain (p < 0.001, η_p_² = 0.345), whereas the aperiodic slope showed almost no association with it (standardised slope = 0.018), indicating a broadband increase in aperiodic activity that left the shape of the spectrum unchanged.

The EEG signal also became more regular and more predictable as physiological strain rose, with a decline in its fractal dimension and entropy and an increase in its compressibility (all p ≤ 0.026, η_p_² = 0.064-0.305, Fig. 1D and Table S1). These changes are consistent with the spectral shift towards slower frequencies. A signal dominated by slow components varies more smoothly from one moment to the next, which makes its course easier to anticipate and its patterns more likely to recur. Fuzzy entropy, which grades how closely the patterns of the signal resemble each other (*16*), was the exception and increased with strain (p < 0.001, η_p_² = 0.217). This increase, together with the decline in the other features, may indicate that the signal came to rely on fewer patterns as strain rose, which recurred more often but with more variation from one occurrence to the next.

To test whether this shift towards slower dynamics also extended to functional connectivity, we estimated weighted symbolic mutual information (wSMI; *18*). The models revealed that coupling increased with physiological strain at the two longer lags, which index slower frequencies (τ = 10 and τ = 40, both p ≤ 0.026, ηp² = 0.236 and 0.066), and decreased at the two shorter ones (τ = 4 and τ = 6, both p ≤ 0.037, ηp² = 0.196 and 0.062; Fig. 1D and Table S1). Mutual information between electrodes thus shifted from fast to slow timescales as exhaustion approached, in step with the redistribution of power towards lower frequencies and the increased regularity of the signal.

### Recovery separates physiological strain from time and muscular demand

During the increasing-effort phase, physiological strain, elapsed time and muscular demand rose together, so any of them could account for the associations reported. The protocol included a recovery phase to dissociate them. After exhaustion, muscular demand fell abruptly and remained low, whereas strain declined gradually and time kept advancing. Changes driven by muscle activity should therefore have collapsed with the load, and an artefact accumulating over the recording should have continued to grow. Instead, most EEG features followed strain back towards baseline (Fig. 1D-E and Table S1). The direction of the effects during increasing effort points the same way. Muscle activity adds power at high frequencies (*19, 20*), however we found that beta power fell and the EEG signal became more predictable and more regular as strain rose. This dissociation is difficult to reconcile with muscle activity or with an artefact building up over time, and supports a reversible shift in neural regime that tracks physiological strain.

### Neural dynamics distinguish the approach to exhaustion

Neural dynamics tracked physiological strain but also marked each participant’s proximity to exhaustion. To establish this, we divided each participant’s increasing-effort phase into 10-s segments and trained a participant-specific gradient-boosted classifier (XGBoost; *21*) that used the EEG features to distinguish the final 5 minutes before exhaustion from all earlier segments (Fig. 2A-B). Discrimination was near-perfect in every participant (AUROC = 0.989 ± 0.007, mean ± SD across participants), and precision–recall analysis confirmed that it was not driven by the predominance of earlier segments (AUPRC = 0.966 ± 0.020; Fig. 2C). Performance was also robust to segment duration (AUROC > 0.91 at every duration tested; Fig. 2D and Table S2). We next confirmed that this high performance was not inflated by the temporal autocorrelation between adjacent segments. With a blocked cross-validation (*22*), in which each test segment was held out together with its neighbours, discrimination remained high (AUROC = 0.942 ± 0.045). Nevertheless, segments at the edges of each held-out block still had neighbours in the training set, so we also removed the training segments closest to each block, within a gap of increasing width. Performance was unchanged with a 10-s gap (AUROC = 0.949 ± 0.037) and decreased progressively as the gap widened to 40 s (AUROC = 0.911 ± 0.059; Fig. 2E and Table S9).

**Figure 2.**
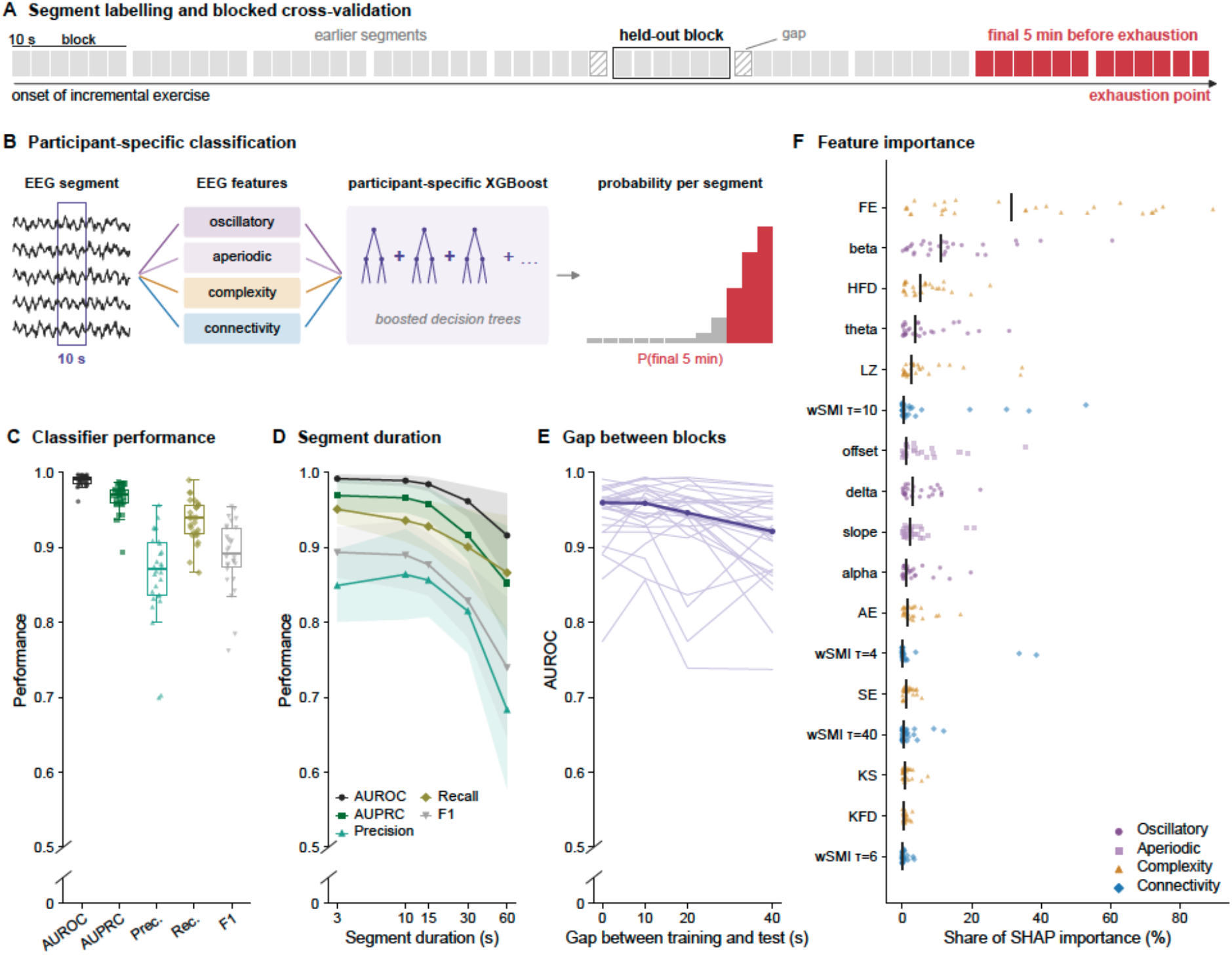
Neural dynamics identify the final minutes before exhaustion. (A) The increasing-effort phase of each participant was divided into consecutive 10-s segments, labelled as belonging to the final 5 min before exhaustion (red) or to earlier segments (grey). For the blocked cross-validation, segments were grouped into contiguous temporal blocks and the held-out block (outlined) was rotated across the recording. A gap of discarded segments on each side of the held-out block (hatched) kept neighbouring segments out of both the training and test sets. (B) Oscillatory, aperiodic, complexity and connectivity features were computed for every segment and entered a gradient-boosted classifier (XGBoost) trained separately for each participant, which returned the probability that each segment belonged to the final 5 min. (C) Classification performance per participant (n = 28) with 10-s segments under repeated stratified k-fold cross-validation. Points are participants and boxes show the median and interquartile range. (D) Performance across segment durations under the same validation (mean ± SD across participants). Each metric keeps its colour and marker in C and D. (E) AUROC under blocked cross-validation as a function of the gap between training and test blocks, for each participant (thin lines) and their median (thick line). (F) Share of SHAP importance of each EEG feature under blocked cross-validation. Points are participants, with colour and marker indicating the feature family, and black lines mark the median. The y axes in C, D and E are truncated. EEG features discriminated the final 5 min before exhaustion in every participant and across segment durations. Discrimination remained high when temporally adjacent segments were kept apart, and declined gradually as the gap widened to 40 s. Classification relied on features from several families, led by fuzzy entropy and followed by beta-band power.

To determine which EEG features supported this discrimination, we computed Shapley additive explanations (SHAP; *23*). Classification relied mainly on signal complexity and oscillatory power features, whereas functional connectivity contributed little at any timescale (Fig. 2F). Fuzzy entropy (33.8% of total importance, mean across participants) and beta power (13.9%) made the largest contributions. The relative weight of each feature, however, varied markedly between participants. Together, these results point to a shared neural signature of the approach tuned to each participant through a different weighting of its components.

We then tested whether neural dynamics carried graded information about proximity to exhaustion, beyond separating the final 5 minutes from the rest of the test, a distinction that treats those minutes as a single state and cannot indicate how close a participant is to stopping. Participant-specific ridge-penalised Cox proportional hazards models (*24*), which rank every segment by its closeness to the end of the effort test (Fig. 3A), reliably reproduced the actual temporal order of segments (C-index = 0.897 ± 0.058; Fig. 3B and Table S3). Because segments from the beginning and the end of the test are easy to tell apart, we restricted concordance to pairs within the final quarter, where successive states are physiologically similar. EEG concordance remained well above chance in this demanding range (C-index = 0.736 ± 0.105), extending this graded information to the final approach to exhaustion. To confirm that these concordance values were not inflated by evaluating each model on the segments used to fit it, we recomputed them out of sample with the blocked cross-validation applied to the classifier, discarding the training segments closest to each held-out block within a gap of increasing width. Concordance was preserved across all gaps, both for the whole test (C-index = 0.875 ± 0.067 without a gap and 0.865 ± 0.072 with a 40-s gap) and for its final quarter (0.678 ± 0.144 and 0.642 ± 0.175; Table S10). The graded information about proximity to exhaustion did not then arise from an artefact of in-sample fitting or from temporal autocorrelation between adjacent segments.

**Figure 3.**
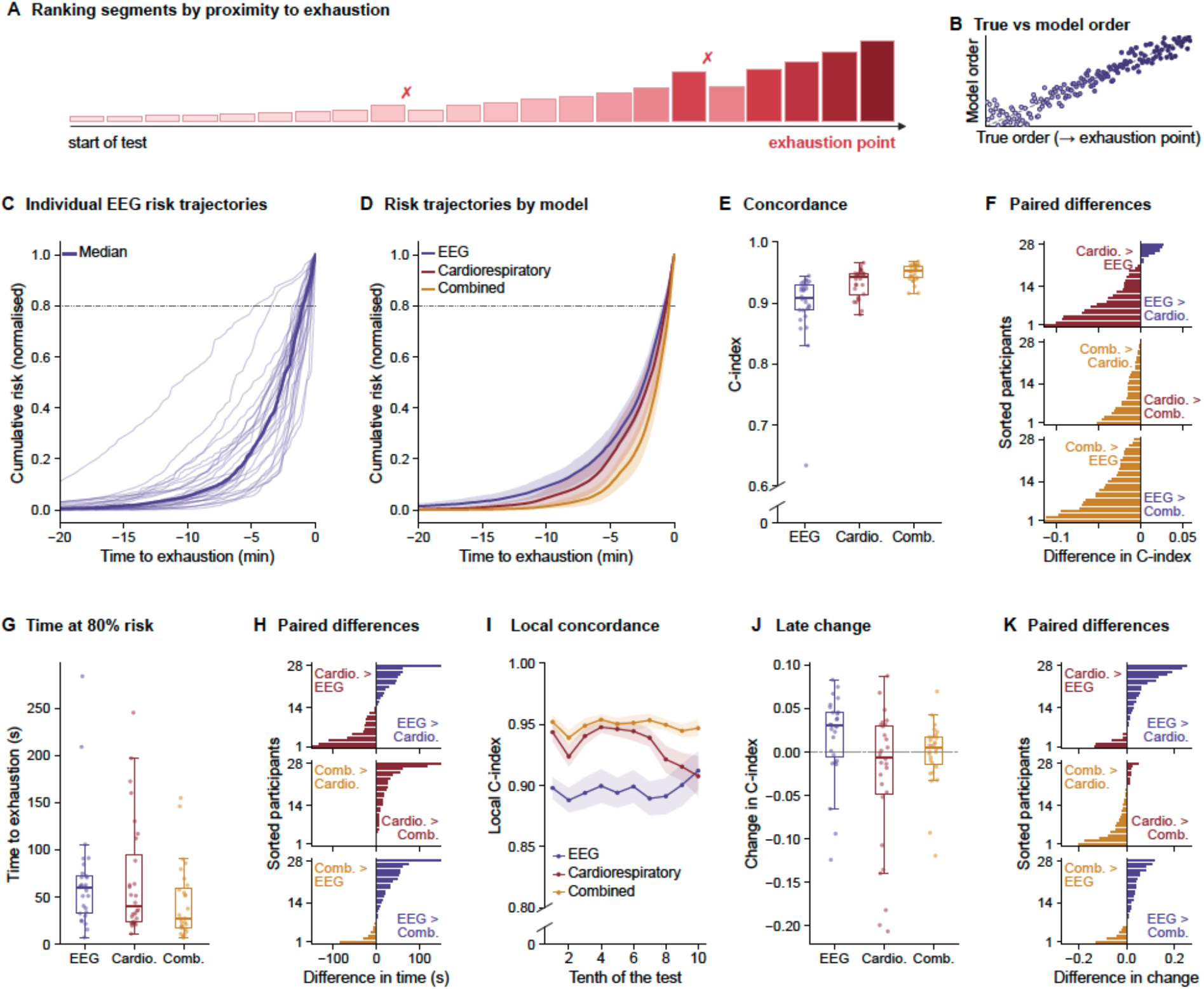
Neural dynamics track proximity to exhaustion and complement cardiorespiratory measures. (A) Participant-specific Cox models assigned a risk score to every 10-s segment of the increasing-effort phase and ranked the segments by their proximity to exhaustion. Concordance (C-index) is the proportion of segment pairs ranked in their true temporal order, and crosses mark pairs ranked in the wrong order. (B) True versus model-assigned order of segments for a representative participant (EEG model). The dashed line marks perfect ranking. (C) Cumulative risk trajectories of the EEG model, normalised to their final value, for each participant (thin lines) and their median (thick line). (D) Mean trajectories of the EEG, cardiorespiratory (Cardio.) and combined (Comb.) models, with shaded 95% bootstrap confidence intervals of the mean. Participants whose test lasted less than 20 min contribute from the start of their test. The dotted line in C and D marks 80% of the final cumulative risk. (E) Concordance of each model per participant across the whole test. (F) Difference in concordance between each pair of models for every participant, sorted by size. Each bar takes the colour of the model with the higher value. (G) Time remaining until exhaustion when the cumulative risk of each model reached 80% of its final value. (H) Difference in that time between each pair of models, displayed as in F. (I) Local concordance along successive tenths of the test (mean ± SEM). (J) Change in concordance over the final tenth of the test relative to the preceding nine, for each model. (K) Difference in that change between each pair of models, displayed as in F. Points are participants, and boxes show the median and interquartile range. The y axes in E and I are truncated. All three models ranked segments by their proximity to exhaustion. Concordance was highest for the combined model, which exceeded the cardiorespiratory model in 27 of 28 participants and the EEG model in all of them. Risk accumulated late in the test and reached 80% of its final value at a median of 60 s (EEG), 40 s (cardiorespiratory) and 27 s (combined) before exhaustion. Over the final tenth of the test, cardiorespiratory concordance declined whereas EEG concordance was maintained in most participants.

To trace how this information built up over the test, we cumulatively summed and normalised the segment-level risk scores of each participant’s Cox model, obtaining an individual trajectory of model-derived proximity to exhaustion (Fig. 3C). Trajectories remained low for most of the test and rose steeply over its final minutes. To quantify this rise, we measured the time remaining until exhaustion once most of the risk had accumulated, taking 80% of the final cumulative risk as the reference level (dotted line in Fig. 3C). This level was reached at a median of 60 s (interquartile range 33–73 s) before exhaustion and within the final 2 minutes in 26 of 28 participants. Results were similar at 70% and 90% (Table S11). The time at the 80% level was only weakly related to total exercise duration (r = 0.22), which varied widely across participants (26.2 ± 6.2 min). The risk estimated from neural dynamics alone was thus concentrated in the final one to two minutes before exhaustion in nearly every participant.

### Neural dynamics complement cardiorespiratory markers near exhaustion

We next benchmarked neural dynamics against cardiorespiratory measures, the standard markers of physiological strain. Applied to the cardiorespiratory measures, the same participant-specific Cox framework achieved higher concordance than the EEG model (C-index = 0.933 ± 0.023; Fig. 3E), exceeding it in 21 of 28 participants (Fig. 3F). Cumulative risk in the cardiorespiratory model reached 80% of its final value at a median of 40 s (interquartile range 23–94 s) before exhaustion, a time that did not differ from that of the EEG model (Wilcoxon signed-rank test, p = 0.90; Fig. 3G and H). Neural dynamics alone tracked proximity to exhaustion almost as accurately as cardiorespiratory measures.

We then tested whether neural dynamics added information to that already captured by cardiorespiratory measures. The combined EEG and cardiorespiratory model achieved the highest overall concordance (C-index = 0.949 ± 0.014; Fig. 3E), exceeding the cardiorespiratory model in 27 of 28 participants and the EEG model in all 28 (Fig. 3F). To separate a genuine EEG contribution from the greater flexibility of a larger model, we compared it with null models in which the EEG features were shuffled in time while the cardiorespiratory measures and model structure were preserved. Temporally aligned EEG improved concordance over this null (ΔC-index = 0.016; Stouffer Z = 17.1, p < 0.001), and this gain held at every segment duration examined (Stouffer Z = 6.0 – 17.6; Table S8). This indicates that neural dynamics carried temporally organised information about proximity to exhaustion that complemented cardiorespiratory measures.

Critically, EEG added most information when the models had to distinguish between successive states close to exhaustion. When concordance was restricted to pairs of segments within the final quarter, temporally aligned EEG produced a gain six times larger than that observed across the complete test (ΔC-index = 0.099 and 0.016, respectively; Table S8). The divergence became clearest over the final tenth of the increasing-effort phase, when cardiorespiratory concordance declined, EEG concordance did not, and the combined model remained stable (change in C-index = −0.028 ± 0.078, +0.017 ± 0.047 and −0.002 ± 0.036, respectively; Fig. 3I and J). This change differed between the three models (Friedman test, χ²(2) = 16.3, p < 0.001). EEG concordance was better preserved than cardiorespiratory concordance in 23 of 28 participants (p = 0.009) and than combined concordance in 21 of 28 (p = 0.034; Holm-corrected Wilcoxon signed-rank tests, Fig. 3K). The decline in cardiorespiratory concordance coincided with a loss of monotonicity in the cardiorespiratory signals near the end of the test, when heart rate plateaued and oxygen uptake and power output fell (*15, 25*), precisely when neural dynamics provided their greatest complementary value.

Combining both modalities concentrated the estimated risk closer to exhaustion. The time at which cumulative risk reached 80% of its final value differed between the three models (Friedman test, χ²(2) = 19.8, p < 0.001). In the combined model this level was reached at a median of 27 s (interquartile range 17–59 s) before exhaustion, later than in the cardiorespiratory model in 26 of 28 participants (p < 0.001) and later than in the EEG model in 21 of 28 (p = 0.012; Holm-corrected Wilcoxon signed-rank tests, Fig. 3G and H). These times also varied less across participants in the combined model than in the cardiorespiratory model (paired permutation test on absolute deviations from the median, p = 0.003). The differences from the cardiorespiratory model held at the 70% and 90% risk levels (Table S11).

We finally tested whether the EEG and cardiorespiratory models generalised across participants. In a leave-one-participant-out scheme, trained on 27 participants and tested on the remaining one, the cardiorespiratory model transferred almost intact (C-index = 0.916 ± 0.059). The EEG model trained on other participants still ranked segments above chance (C-index = 0.781 ± 0.081), but it no longer produced the steep rise in risk before exhaustion that the participant-specific models showed. Its partial transfer indicates that the reorganisation was shared across participants but the signal of proximity to exhaustion followed a time course specific to each individual.

## DISCUSSION

Physical exhaustion has long been described through the cardiorespiratory and metabolic cost of exercise. Here we show that neural dynamics also carry a reliable signature of its approach, detectable in scalp EEG, that allows it to be anticipated. As effort rises, brain activity becomes progressively slower and more predictable, and is coupled over longer timescales. Time-to-event models based on these features tracked the final minutes before participants stopped exercising. Their performance approached that of classical cardiorespiratory measures, and combining both sources estimated proximity to exhaustion even more accurately. When trained across participants, the same models showed that this signature is common to all individuals, although its time course differs from one person to another.

As exhaustion approached, the aperiodic, non-rhythmic component of the EEG signal rose across the whole spectrum with almost no change in its slope. By contrast, rhythmic power above that background increased only in the delta band and declined in faster frequency bands (i.e., theta, alpha and beta). These findings revise the existing literature on brain activity during high-intensity exercise, which has systematically reported an increase in power across the whole EEG spectrum (e.g., *10, 26*). Our findings confirm this broadband rise in EEG power with effort but point to the non-rhythmic activity of neuronal populations as its source (*27, 28*). However, a broadband rise in power could also be expected from the muscle contamination of the EEG, which would increase monotonically with effort. In that case, muscle contamination would add power predominantly at fast frequencies (i.e., above 20 Hz; *19, 20, 29*), but the only increase in rhythmic power reported here unfolded in the slowest frequency band. Further, an increase in power at high frequencies would have flattened the slope of the spectrum. Yet the slope remained unchanged. Muscle contamination is therefore unlikely to account for either of the two features that define these results, the preserved slope of the spectrum and the specific pattern of changes in rhythmic power as physical effort increased.

In parallel, neural dynamics became more regular and more predictable as exhaustion approached, and functional coupling shifted towards longer timescales, in line with the lengthening of intrinsic neural timescales when the brain moves from rest into active behavioural states (*30, 31*). A more regular signal cycles through a smaller repertoire of patterns, which reduces its capacity to carry information (*32*). This pattern of change is characteristic of complex systems that are gradually pushed towards a critical transition, in which a small further increase in demand produces an abrupt shift in their behaviour (*13*). The approach to exhaustion has this structure, as effort builds up over many minutes but ends abruptly when the participant reduces their effort deployment, eventually stopping the exercise. Exhaustion is not the only physiological state reached in this way. In the minutes that precede sleep onset, EEG dynamics slow down and become more predictable (*14*). The loss of consciousness under propofol is also accompanied by a fall in signal complexity (*33*) and a selective gain in delta power (*28*). The approach to exhaustion shares some of these features, which suggests that exhaustion may be the end point of a comparable transition that develops as effort rises. In each of these cases, a sustained demand (e.g., sleep pressure, an anaesthetic or physical effort) shifts the system away from the dynamics of wakeful rest, and removing the demand does not bring it back along the same path. Our findings during the recovery phase are consistent with this view. After exercise stopped, neural dynamics returned towards their resting values along a different path, as the same level of physiological strain was accompanied by different dynamics than during increasing effort. Physical effort to exhaustion thus offers a way to push a physiological system far from rest and to observe how it returns, which makes the hysteresis of neural dynamics directly measurable.

Neural dynamics traced this transition and revealed how close each participant was to its end. This information was spread across a multidimensional profile of neural changes, with signal complexity and oscillatory power carrying most of it. The profile was shared across individuals, but the contribution of each feature differed from one participant to the next. Indeed, models trained on other participants still tracked proximity to exhaustion above chance, which reveals a common pattern, whereas their loss of precision shows that each individual expresses it with its own weighting of features and time course. The approach to exhaustion would combine a common neural signature with a configuration specific to each individual. EEG features tracked proximity to exhaustion with an accuracy comparable to that of cardiorespiratory measures. Most of the information added by the EEG came at the end of the test, where cardiorespiratory measures became less informative, presumably due to the well-described plateau of cardiorespiratory signals close to maximal effort observed in some individuals (*15, 25*). Neural dynamics contribute most when cardiorespiratory signals stop changing monotonically, which is also when an estimate of proximity to exhaustion is most needed. The combined model (EEG and cardiorespiratory features) placed the risk of exhaustion closer to the moment at which it occurred and reduced the variation between participants than either model alone. The complementarity between neural and cardiorespiratory measures opens the way to training existing models for the prediction of exhaustion with personalised EEG data, collected from each individual over repeated sessions. Low-cost wearable devices could then bring these personalised models into everyday use.

These findings nevertheless call for caution, because they come from a single incremental test, in which workload, physiological strain and elapsed time rise together. The variation in workload and duration between participants, across which anticipation held, reduces this concern but cannot remove it, as the three factors still rose together within each test. The cumulative risk of a Cox model, moreover, rises towards the end of the test by construction and is scaled to a final moment that is only known once the test has ended. Stringent preprocessing and a validation that kept neighbouring segments apart limit the influence of noise and temporal autocorrelation, but cannot exclude that noise accumulating over the recording accounts for part of these results. Exercise protocols in which strain and time do not rise together, and repeated sessions in the same individuals, are needed to rule out these explanations and to establish whether an individual calibration holds over time.

Physical exhaustion should therefore be understood not only as a cardiorespiratory and metabolic limit, but also as the end point of a transition in neural dynamics that unfolds within each individual. Tracking this transition at the neural level opens new avenues for anticipating exhaustion when cardiorespiratory markers lose sensitivity, and for addressing the mechanisms that determine the limits of exercise tolerance.

## METHODS

### Participants

Thirty-two trained amateur cyclists were selected to take part in this experiment. Four were excluded due to data acquisition issues, leaving 28 participants (4 women; age 28.2 ± 11.0 years, range 18–52). Participants were healthy individuals with no history of cardiovascular, metabolic, neurologic, or psychiatric disorders. They all reported normal or corrected-to-normal vision and hearing. All participants trained on the bike at least three times a week during the standard training season. Participants were instructed not to consume coffee the day of the experiment and not to train in the 24h prior to the testing session.

Participants were recruited by word of mouth and gave their written informed consent prior to taking part in the experiment and received a compensation of €60. The study was approved by the University of Granada’s Ethics Committee (978/CEIH/2019). The general methodology of the study and preliminary analyses were preregistered on the Open Science Framework before data collection (*34*). The relevant deviations from the initial preregistration are listed in Table S13.

### Procedure

Participants were prepared for data collection in a Faraday cage, where they were fitted with a 64-channel electroencephalography (EEG) system (actiCAP, Brain Products GmbH, Germany) and a gas analyser mask (Geratherm Respiratory GmbH, Germany) to measure the volume of gas exchange. Written instructions about the session were provided during the participants’ preparation process.

The experimental session comprised seven consecutive periods designed to track EEG and cardiorespiratory activity from rest to maximal physical exertion and back (Fig. 1A). First, a 5-min eyes-closed resting state (RS0) was recorded while participants sat comfortably on a chair, followed by a 5-min eyes-open resting state (RS1). Participants were then fitted with a chest-strap heart rate (HR) monitor (HRM-Pro, Garmin Ltd., USA) and moved to an indoor cycle ergometer (SRM GmbH, Germany) equipped with Assioma PRO RL power meter pedals (Favero Electronics srl, Italy). Subsequently, participants cycled at low intensity for a 5-min warm-up. They then began an incremental protocol starting at 120 W for men and 100 W for women, with increments of 30 W every 5 min. The load of the ergometer was controlled with Gobat software (Magnetic Days, Italy). Participants were asked to choose their preferred cadence during the warm-up and to keep it constant throughout the protocol, always above 60 rpm, and were encouraged to continue pedalling until exhaustion. The incremental protocol ended when participants stopped pedalling abruptly or could no longer maintain a cadence above 60 rpm, at which point the resistance of the ergometer was lowered. This time point marked the end of the incremental protocol in all analyses and was the event from which proximity to exhaustion was computed. Immediately after the incremental protocol, participants cycled at low intensity for 5 min (recovery) and then completed a 5-min eyes-open resting state (RS2) and a 5-min eyes-closed resting state (RS3), seated on a comfortable chair.

A modified version of the auditory global–local paradigm (35) ran in the background throughout the session. It consisted of a passive task, in which pairs of brief tones were presented continuously through in-ear headphones. Participants only had to press a button on the handlebar when one of the rare target sounds was presented (two targets per 5-min block) as catch trials. These data address secondary hypotheses unrelated to the main aim of the present study and have not been examined yet.

### EEG recording and preprocessing

Continuous 64-channel EEG signals were acquired at a 1,000 Hz sampling rate using an actiCHamp amplifier and Brain Vision software (Brain Products GmbH, Munich, Germany). Electrodes were positioned according to the extended 10–20 system, referenced online to FCz, with the ground electrode at the centre of the forehead. Electrode impedances were kept below 10 kΩ throughout the session. Offline preprocessing was performed in MATLAB with custom scripts and the EEGLAB (36) and FieldTrip (37) toolboxes. The continuous recording was first high-pass filtered at 0.5 Hz, low-pass filtered at 126 Hz, notch filtered at 50 and 100 Hz and downsampled to 500 Hz. Physiological artifacts were then isolated with an independent component analysis (ICA, runica algorithm), and the components corresponding to blinks, eye movements and, when present, the cardiac artifact were identified by visual inspection and removed (median of 4 components per participant, range 1–7). The data were then downsampled to 256 Hz and band-pass filtered between 1 and 40 Hz to remove most of the remaining noise from muscle activity, movement, sweating and respiration. The continuous recording was divided into the experimental periods and segmented into consecutive 3-s epochs. Epochs were flagged as noisy by an automated procedure if their spectral power deviated from the mean by more than ±50 dB in the 0–2 Hz range (to target ocular drift) or by more than +25 or −100 dB in the 20–40 Hz range (to target muscle activity). Noisy channels were interpolated with spherical splines (one channel in one participant). The online reference electrode (FCz) was then restored and the signal was re-referenced to the average of all electrodes. To preserve the temporal continuity required by the research question, noisy epochs were not removed but replaced by the nearest preceding clean epoch.

### Cardiorespiratory and power output measures

Concurrently to the cycling, power output and cadence data were captured using a PowerControl 8 (SRM GmbH, Germany) paired with the bike pedals and the HR monitor (1Hz sampling rate). Gas exchange was measured as the volume of oxygen (VO_2_), VO_2_ over bodyweight (VO_2_/kg), the volume of carbon dioxide (VCO_2_) and the respiratory exchange ratio (RER), with a sampling rate corresponding to one sample per respiration. The stopping of the recording of the cardiorespiratory and power output measures during the incremental test was done manually.

The recordings of VO_2_, VO_2_/kg, VCO_2_, RER, HR and power output were synchronised with the EEG recording. All measurements were linearly interpolated to 1 Hz and then averaged within the 3-s windows of the EEG epochs. Given occasional measurement errors of the power meter pedals, implausible values (negative power or cadence, power above 600 W and cadence above 120 rpm) were removed and linearly interpolated.

### Neural measures

#### Spectral analysis

The analysis was performed on the consecutive 3-s EEG epochs obtained during preprocessing. We applied the Fit Oscillations and One Over F (FOOOF; *17*) method using the specparam algorithm implemented in FieldTrip to obtain the aperiodic and corrected oscillatory components of the signal. Spectral estimates were computed in the 1–40 Hz frequency range using a multitaper fast Fourier transform approach (mtmfft), with zero-padding to 4 s and a spectral smoothing of 2 Hz. The aperiodic component was fitted independently for each epoch and channel. For every epoch and channel, FieldTrip’s FOOOF implementation was used to estimate the fitted aperiodic spectrum and its intercept and exponent. The aperiodic slope was then defined as the negative of the FOOOF exponent. The aperiodic-adjusted oscillatory spectrum was calculated as 10·log_10_(P_original_/P_aperiodic_), yielding an oscillatory-to-aperiodic power ratio expressed in dB. For each participant and block, the trial-wise aperiodic parameters and oscillatory spectra were extracted. Oscillatory power was extracted and averaged within delta (1–4 Hz), theta (4–8 Hz), alpha (8–13 Hz), and beta (13–30 Hz) frequency bands. Finally, values within each individual spectral feature were averaged across channels.

#### Complexity measures

EEG complexity was quantified on the consecutive 3-s epochs of the whole recording, separately for each participant, channel and epoch. Seven measures were computed, Katz fractal dimension (KFD), Higuchi fractal dimension (HFD), Kolmogorov-Sinai entropy (KS), sample entropy (SE), approximate entropy (AE), fuzzy entropy (FE) and Lempel–Ziv complexity (LZ), chosen to capture both the predictability and the regularity of the signal (16). SE, AE, FE and KS were computed with the EntropyHub toolbox (v2.0), with a time delay of 1 and a tolerance of 0.2 times the standard deviation of the epoch. The embedding dimension was 1 for SE and AE and 2 for FE and KS, and FE used the default exponential membership function (r = 0.2, n = 2). KS was estimated as the correlation (K2) entropy. HFD was computed with k_max_ = 60, and LZ with the LZ78 algorithm on the signal after detrending and binarising it around zero. Each complexity value was normalised with a shuffled surrogate of the same epoch. The time series was randomly permuted, which preserves its amplitude distribution but destroys its temporal order, the measure was recomputed on the shuffled series, and the final value was the ratio between the original and the shuffled estimates. This normalisation was applied independently for each participant, period, channel, epoch and measure. The normalised values were then averaged across channels, yielding one whole-scalp value per epoch and measure.

#### Weighted symbolic mutual information

We quantified information sharing between electrodes using weighted symbolic mutual information (wSMI; 11, 18, 38) on continuous 3-s epochs. wSMI estimates statistical dependencies between symbolic representations of EEG signals and is sensitive to nonlinear coupling, while reducing spurious correlations that may arise from common sources or volume conduction. wSMI was computed separately for each consecutive epoch, thus obtaining a continuous measure of wSMI across time. Before symbolisation, each epoch was low-pass filtered at fs/(k·τ) with a zero-phase sixth-order Butterworth filter to prevent aliasing (21.3, 14.2, 8.5 and 2.1 Hz for τ = 4, 6, 10 and 40, respectively). EEG signals were then transformed into sequences of discrete symbols using a kernel size of k = 3. wSMI was computed for four time lags, τ = 4, 6, 10 and 40 samples (15.6, 23.4, 39.1 and 156.3 ms at 256 Hz), selected to probe information sharing across a range of temporal scales, so that τ = 4 probes the fastest interactions and τ = 40 the slowest. The analysis was performed over the full epoch duration, using all available samples in each epoch, and wSMI was calculated between all pairs of electrodes. The joint probability distribution of symbol pairs was weighted with the standard wSMI weights, which discard identical and opposite symbol pairs, since both are likely to arise from common sources or from the two poles of a single dipole rather than from genuine information sharing. For each τ, wSMI was finally averaged across all 2,016 electrode pairs, yielding a single global connectivity estimate per epoch.

### Neural dynamics and physiological strain

The epoch-wise neural and cardiorespiratory measures were first averaged within each experimental period from RS1 to RS2, and within consecutive sub-blocks of the incremental protocol, so that each participant contributed one value per block and measure. Before averaging, the cardiorespiratory measures (VO_2_, VO_2_/kg, VCO_2_, RER and HR) were z-scored within each participant to make them comparable across participants. Because substantial collinearity was expected among the cardiorespiratory measures, we computed pairwise Kendall’s τ correlations between them within each participant, across all blocks, and averaged the resulting correlation matrices across participants. Kendall’s τ is a rank-based measure that is robust to outliers and does not assume a linear relationship. We then performed principal component analysis (PCA) on the block-averaged, standardised cardiorespiratory measures of all participants to derive a single composite index of physiological strain. Only the first component was retained for the subsequent analyses (see Statistical analysis).

### Classification of the final minutes before exhaustion

#### EEG segmentation and feature extraction

The incremental exercise recording was divided into non-overlapping segments of fixed duration to enable a time-resolved, segment-level characterization of brain dynamics. Five segment lengths were evaluated (3, 10, 15, 30 and 60 s). For each participant, segments were extracted from the onset of the incremental protocol until the participant ceased exercising, yielding a variable number of segments per participant depending on individual exercise tolerance. Segments were labeled according to their temporal proximity to exhaustion. Those whose end fell within the final five minutes preceding exercise cessation were assigned to the near-exhaustion class (label = 1), while all earlier segments were assigned to the far-from-exhaustion class (label = 0). This binary labeling scheme resulted in a class-imbalanced dataset, with the far-from-exhaustion class substantially outnumbering the near-exhaustion class for most participants.

#### Binary classification

A participant-specific binary classification approach was adopted to evaluate the discriminability of the extracted EEG features with respect to proximity to exhaustion. It was done independently for each participant to avoid confounds from inter-individual differences in neural activity levels and exercise capacity, using all available segments with labels assigned as described in the previous section (near-exhaustion: label = 1; far-from-exhaustion: label = 0). Each segment length (3, 10, 15, 30, and 60 s) was analysed separately.

Prior to classification, all features were standardised to zero mean and unit variance using a StandardScaler fitted exclusively on the training fold and applied to the test fold without refitting. To address the class imbalance resulting from the binary labelling scheme (whereby the far-from-exhaustion class substantially outnumbers the near-exhaustion class) the Synthetic Minority Over-sampling Technique (SMOTE; *39*) was applied to the standardised training set. The number of nearest neighbours for synthetic sample generation was set adaptively as *k* = max(1, min(5, *n* − 1)), where *n* is the number of minority-class training samples in that fold, preventing SMOTE failures when very few minority samples are available. Classification was performed using an Extreme Gradient Boosting classifier (XGBoost; *21*). Hyperparameters were optimised within each fold via a randomised search (*n* = 10 parameter combinations, 3-fold inner cross-validation, F1 score as optimisation criterion) over ranges of the number of estimators {100, 200, 500, 1000}, maximum tree depth {3, 5, 7, 9, 12}, learning rate {0.001, 0.01, 0.05, 0.1, 0.2}, subsampling ratio {0.5, 0.7, 0.9, 1.0}, column subsampling ratio {0.5, 0.7, 0.9, 1.0}, minimum loss reduction γ {0, 0.1, 0.5, 1}, and minimum child weight {1, 3, 5, 7}.

### Proximity to exhaustion

To model the temporal dynamics of approaching exhaustion, a Cox proportional hazards (PH) model (*24*) was fitted to each participant’s feature time series. The same survival framework was applied to three feature sets (the EEG features, the cardiorespiratory features and their combination by early fusion). This framing treats exercise cessation as the event of interest and positions each segment as an observation with a known time-to-event, given by the number of segments remaining until exhaustion. Specifically, each segment t was assigned a duration value of n - t -1, where n is the total number of segments for that participant, such that the final segment before exercise cessation received a duration of zero. All segments were assigned an event indicator of 1, as exhaustion was observed in all participants.

Prior to model fitting, EEG features were standardised to zero mean and unit variance. To remove the influence of artifact-driven outliers (isolated segments with extreme spectral or complexity values attributable to electrode noise or movement) standardised features were winsorised by clipping values exceeding ±5 standard deviations. Winsorised features were then submitted to PCA. The number of components retained was determined adaptively per participant by selecting the minimum number of components required to explain 70% of the total variance. This threshold was chosen to balance representational fidelity with model parsimony, ensuring a sufficient ratio of observations to predictors for the penalised Cox model given the variable number of segments per participant. Retaining fewer components also reduces the risk of overfitting inherent in high-dimensional Cox regression, even under ridge penalisation. The resulting concordance index values confirmed that the retained components captured the temporally relevant structure of the EEG signal.

The cardiorespiratory model was fitted to the five cardiorespiratory variables (VO_2_, VO_2_/kg, VCO_2_, RER and HR), which were standardised and winsorised in the same manner but, given their low dimensionality, were entered directly into the model without PCA. The combined model was constructed by early fusion. For each participant, the retained EEG principal components were concatenated with the standardised cardiorespiratory features before fitting a single Cox model, so that both feature blocks entered the model on a comparable scale and neither modality dominated by virtue of its dimensionality. A ridge-penalised Cox PH model (penaliser = 0.1) was fitted independently for each participant using the lifelines library (40). From the fitted model, the instantaneous risk of exhaustion at each segment was estimated as one minus the predicted survival probability. A cumulative probability curve was then constructed as the normalised cumulative sum of these instantaneous risks, yielding a monotonically increasing trajectory bounded between 0 and 1 that reflects the accumulated evidence of impending exhaustion over the course of the exercise protocol.

The temporal anticipation afforded by each model was defined as the time remaining before exercise cessation at which the cumulative-risk curve reached 80% of its final value. For each participant, the curve was expressed as a function of the fraction of the test completed, and the fraction f_80_ at which it first reached 0.8 was obtained by linear interpolation. Anticipation was then computed as (1 − f_80_) · n·W, where n is the number of segments and W the segment duration in seconds. This criterion requires no smoothing, no search window and no tuning parameter, and it yields a valid estimate in every participant. To verify that conclusions did not depend on the chosen level, anticipation was also computed at 70% and 90% of the final value (Table S11). Because anticipation times were right-skewed across participants, they are summarised as the median and interquartile range (IQR). The proportion of participants warned within the final two minutes is reported as an indicator of practical usefulness.

### Statistical analysis

#### Physiological strain and neural dynamics

To test the relationship between physiological strain and neural activity, a piecewise linear model was fitted to each EEG feature, with strain and phase as predictors. Strain was indexed by the first principal component of the cardiorespiratory measures (PC1), which captured their shared variance, and the EEG features were z-scored to make them comparable. The piecewise model followed the experimental protocol, which was divided at exhaustion into an increasing-effort phase (RS1, warm-up and incremental protocol) and a recovery phase (low-intensity cycling and RS2). The eyes-closed resting states (RS0 and RS3) were not included. For each EEG feature, a reduced model including participant and phase as predictors was compared with a full model that additionally included strain and its interaction with phase. Participant was included as a fixed effect to account for between-participant differences in baseline neural activity. The comparison between the two models yielded a partial F-statistic testing whether strain explained additional variance in the feature, either as a main effect or as a phase-dependent effect, and effect sizes were expressed as partial eta squared (η_p_²). To correct for multiple comparisons within each family of EEG features (oscillatory, aperiodic, complexity and wSMI), statistical significance was assessed with a permutation approach. In each permutation, the values of physiological strain were randomly shuffled within participant and phase, which preserves the participant-specific structure, the phase structure and the phase-specific distribution of strain while disrupting its association with neural activity. The partial F-statistic was recomputed for every EEG feature in each permuted dataset, and the maximum F-statistic across the features of each family was retained. This procedure was repeated 10,000 times to generate family-wise empirical null distributions. The F-statistic of each feature in the original data was then compared with the null distribution of its family, and its p-value was computed as the proportion of permutations, counting the observed one, whose maximum F-statistic equalled or exceeded it. Effects were considered significant at α = 0.05.

#### Classification performance

Classification performance was estimated with a repeated stratified k-fold cross-validation (k = 5, 2 repetitions, yielding 10 test folds per participant), which preserves the class ratio across folds. Rather than applying a fixed decision threshold, the threshold was optimised on the training partition of each fold, as the value between 0.1 and 0.9 (in steps of 0.01) that maximised the F1 score, and was then applied unchanged to the held-out fold, so that threshold-dependent metrics were not optimistically biased by the test data. Performance was quantified with the area under the receiver operating characteristic curve (AUROC), the area under the precision–recall curve (AUPRC), and precision, recall and F1 score at the optimised threshold. To account for the variability introduced by SMOTE and the randomised hyperparameter search, the whole cross-validation was repeated five times with different random seeds, and the performance of each participant was averaged across the resulting 50 test folds (5 repetitions × 10 folds).

#### Feature importance

The contribution of each EEG feature to the classification was estimated with SHapley Additive exPlanations (SHAP; 23) computed on out-of-fold predictions. For each participant, the ordered sequence of segments was divided into ten contiguous blocks (blocked cross-validation with no gap, see Control and robustness analyses). An XGBoost classifier with fixed hyperparameters was trained on nine blocks, and TreeSHAP values were computed for the segments of the held-out block, so that every SHAP value was obtained on data not used to fit the model. For each participant, the mean absolute SHAP value of each feature was computed across all segments and expressed as a percentage of the sum across features, yielding a relative importance profile. Importance was aggregated by feature type, summing over the 64 channels for each of the 13 channel-level features, while each of the four global wSMI features formed a feature type of its own. Group-level importance is reported as the mean of the per-participant profiles, and feature types are ordered by their median across participants, which is robust to the few participants with extreme values. Per-channel importance was obtained in the same way, summing over feature types instead of channels.

#### Survival models and model comparisons

The discrimination of the survival models was assessed with Harrell’s concordance index (C-index; 41), the proportion of segment pairs that a model ranks correctly according to their proximity to exhaustion. A C-index of 0.5 indicates chance-level discrimination and a value of 1 a perfect ranking. In addition to the global C-index, computed over all segment pairs, a final-phase C-index was restricted to pairs in which both segments fell within the final 25% of the test. Segments early in the test are trivially distinguished from those close to exhaustion, so the global index is dominated by easy comparisons, whereas the final-phase index quantifies discrimination among segments close to exhaustion, where prediction matters most. Local concordance was computed in the same way within each tenth of the test (pairs of segments both lying in that tenth). The change in concordance near exhaustion was defined as the C-index over the final tenth minus that over the first 90% of the test, and compared between modalities with a Wilcoxon signed-rank test. Anticipation times of the three models were compared with a Friedman test, followed by pairwise Wilcoxon signed-rank tests with Holm correction. Whether fusion reduced the dispersion of anticipation across participants was tested with a sign-flip permutation test on the per-participant absolute deviations from the cohort median (10,000 permutations). All tests were two-sided unless stated otherwise, with α = 0.05.

### Control and robustness analyses

#### Blocked cross-validation with a gap

To provide a leakage-controlled complement to the repeated stratified k-fold evaluation, in which temporally adjacent segments may fall in both the training and the test fold, the classification was repeated under a blocked cross-validation with a gap (*22*). For each participant, the ordered sequence of segments was divided into ten contiguous blocks. Each block served in turn as the test fold, and the remaining blocks as the training fold. To prevent leakage across fold boundaries, training segments lying within a gap of the test block were removed. Gaps of 0, 10, 20 and 40 s were examined. With a gap of 0 s the folds are simply contiguous and non-overlapping. Class balancing, standardisation and the XGBoost classifier were applied exactly as in the main scheme, and discrimination was quantified on the pooled out-of-fold predictions of each participant (Table S9).

#### Out-of-sample concordance of the survival models

Because the within-participant Cox models were fitted and evaluated on the same segments, we also estimated their concordance out of sample. For each participant, the ordered segments were divided into ten contiguous blocks. Standardisation, winsorisation, PCA and the Cox model were fitted on nine blocks only and used to predict the risk of the held-out block, and the C-index was computed on the pooled out-of-fold predictions. Gaps of 0, 10, 20 and 40 s were removed from the training data on either side of each test block (see Blocked cross-validation with a gap). Results are reported in Table S10.

#### Complementary information of neural and cardiorespiratory features

To verify that the advantage of the combined model did not merely reflect its larger number of free parameters (the combined feature set being the union of the EEG and cardiorespiratory feature sets) we used a permutation test of complementary information. The aim was not to rank the modalities, as cardiorespiratory models are well established, but to test whether the neural features carried information about the approach to exhaustion beyond that already provided by the cardiorespiratory variables, while holding model dimensionality fixed. For each participant and segment length, a reduced Cox model was fitted on a base modality and a full model on the base plus the added modality. The statistic of interest was the C-index of the full model, computed both globally and within the final 25% of the task. A null distribution was generated by randomly permuting the temporal order of the added modality’s feature rows (10,000 permutations) and refitting the full model on each permutation, which preserves the added block’s dimensionality and marginal distribution while destroying its alignment with the progression of the exercise, so that any concordance it contributes under the null arises solely from the additional degrees of freedom. A one-sided p-value was computed as the proportion of permutations whose C-index equalled or exceeded the observed value. Per-participant p-values were combined across the cohort using Stouffer’s method, and the effect size was summarised as the difference between the observed C-index and the mean of the permutation null. The test was applied in both directions (EEG added to cardiorespiratory features, and cardiorespiratory features added to EEG) at every segment length.

#### Between-participant survival analysis (leave-one-participant-out)

To test whether the survival mapping generalised across individuals, the ridge-penalised Cox model was refitted in a leave-one-participant-out (LOPO) scheme. For each held-out participant, the model was trained on the remaining 27 participants (stratified by participant) and evaluated on the held-out one, with all preprocessing (standardisation, winsorisation and, for EEG, the 70%-variance PCA) fitted on the training participants only, so that every estimate was fully out of sample. Transfer was quantified by the C-index on the held-out participant and by the shape of the cumulative-risk trajectory, for both the EEG and the cardiorespiratory feature sets (Fig. S17B, Table S12).

## Data and code availability

The data are available on Zenodo (https://doi.org/10.5281/zenodo.23919810) and the analysis code on OSF (https://doi.org/10.17605/OSF.IO/7ENRG).

## Author contributions

Conceptualization: C.Av., T.A.B., D.S., L.F.C. Methodology: C.Av., C.Al., T.A.B., D.S., L.F.C. Investigation: C.Av., D.S., L.F.C. Formal analysis: C.Av., J.E.A., C.Al., L.F.C. Visualization: C.Av., J.E.A., C.Al., L.F.C. Funding acquisition: C.Av., T.A.B., D.S., L.F.C. Supervision: C.Av., D.S., L.F.C. Writing – original draft: C.Av., J.E.A., D.S., L.F.C. Writing – review & editing: C.Av., J.E.A., C.Al., T.A.B., D.S., L.F.C.

## Use of AI-assisted technologies

The authors used AI-assisted tools to refine the wording of text they drafted and to check it for potential errors, and to help diagnose and fix errors in analysis and plotting code under their supervision. The authors reviewed and verified every AI-assisted output and take full responsibility for the content of the manuscript. No data, image or figure was generated by an AI model.

## REFERENCES

1. D. M. Bramble, D. E. Lieberman, Endurance running and the evolution of *Homo*. Nature 432, 345–352 (2004).

2. D. A. Raichlen, G. E. Alexander, Adaptive capacity: An evolutionary neuroscience model linking exercise, cognition, and brain health. Trends Neurosci. 40, 408–421 (2017).

3. A. D. Craig, How do you feel? Interoception: The sense of the physiological condition of the body. Nat. Rev. Neurosci. 3, 655–666 (2002).

4. J. L. Taylor, M. Amann, J. Duchateau, R. Meeusen, C. L. Rice, Neural contributions to muscle fatigue: From the brain to the muscle and back again. Med. Sci. Sports Exerc. 48, 2294–2306 (2016).

5. M. Hargreaves, L. L. Spriet, Skeletal muscle energy metabolism during exercise. Nat. Metab. 2, 817–828 (2020).

6. B. Kayser, Exercise starts and ends in the brain. Eur. J. Appl. Physiol. 90, 411–419 (2003).

7. C. V. Robertson, F. E. Marino, Prefrontal and motor cortex EEG responses and their relationship to ventilatory thresholds during exhaustive incremental exercise. Eur. J. Appl. Physiol. 115, 1939–1948 (2015).

8. L. F. Ciria, P. Perakakis, A. Luque-Casado, D. Sanabria, Physical exercise increases overall brain oscillatory activity but does not influence inhibitory control in young adults. NeuroImage 181, 203–210 (2018).

9. B. Chen, L. F. Ciria, C. Hu, P. Ch. Ivanov, Ensemble of coupling forms and networks among brain rhythms as function of states and cognition. *Commun*. Biol. 5, 82 (2022).

10. L. Hosang, E. Mouchlianitis, S. M. R. Guérin, C. I. Karageorghis, Effects of exercise on electroencephalography-recorded neural oscillations: A systematic review. Int. Rev. Sport Exerc. Psychol. 17, 926–979 (2024).

11. J. D. Sitt, J.-R. King, I. El Karoui, B. Rohaut, F. Faugeras, A. Gramfort, L. Cohen, M. Sigman, S. Dehaene, L. Naccache, Large scale screening of neural signatures of consciousness in patients in a vegetative or minimally conscious state. Brain 137, 2258–2270 (2014).

12. D. Iannetta, E. C. Inglis, A. T. Mattu, F. Y. Fontana, S. Pogliaghi, D. A. Keir, J. M. Murias, A critical evaluation of current methods for exercise prescription in women and men. Med. Sci. Sports Exerc. 52, 466–473 (2020).

13. M. Scheffer, J. Bascompte, W. A. Brock, V. Brovkin, S. R. Carpenter, V. Dakos, H. Held, E. H. van Nes, M. Rietkerk, G. Sugihara, Early-warning signals for critical transitions. Nature 461, 53–59 (2009).

14. J. Li, A. Ilina, R. Peach, T. Wei, E. Rhodes, V. Jaramillo, I. R. Violante, M. Barahona, D.-J. Dijk, N. Grossman, Falling asleep follows a predictable bifurcation dynamic. Nat. Neurosci. 28, 2515–2525 (2025).

15. M. Niemeyer, R. Knaier, R. Beneke, The oxygen uptake plateau—A critical review of the frequently misunderstood phenomenon. Sports Med. 51, 1815–1834 (2021).

16. Z. J. Lau, T. Pham, S. H. A. Chen, D. Makowski, Brain entropy, fractal dimensions and predictability: A review of complexity measures for EEG in healthy and neuropsychiatric populations. Eur. J. Neurosci. 56, 5047–5069 (2022).

17. T. Donoghue, M. Haller, E. J. Peterson, P. Varma, P. Sebastian, R. Gao, T. Noto, A. H. Lara, J. D. Wallis, R. T. Knight, A. Shestyuk, B. Voytek, Parameterizing neural power spectra into periodic and aperiodic components. Nat. Neurosci. 23, 1655–1665 (2020).

18. J.-R. King, J. D. Sitt, F. Faugeras, B. Rohaut, I. El Karoui, L. Cohen, L. Naccache, S. Dehaene, Information sharing in the brain indexes consciousness in noncommunicative patients. Curr. Biol. 23, 1914–1919 (2013).

19. I. I. Goncharova, D. J. McFarland, T. M. Vaughan, J. R. Wolpaw, EMG contamination of EEG: Spectral and topographical characteristics. Clin. Neurophysiol. 114, 1580–1593 (2003).

20. E. M. Whitham, K. J. Pope, S. P. Fitzgibbon, T. Lewis, C. R. Clark, S. Loveless, M. Broberg, A. Wallace, D. DeLosAngeles, P. Lillie, A. Hardy, R. Fronsko, A. Pulbrook, J. O. Willoughby, Scalp electrical recording during paralysis: Quantitative evidence that EEG frequencies above 20 Hz are contaminated by EMG. Clin. Neurophysiol. 118, 1877–1888 (2007).

21. T. Chen, C. Guestrin, “XGBoost: A scalable tree boosting system” in Proceedings of the 22nd ACM SIGKDD International Conference on Knowledge Discovery and Data Mining (Association for Computing Machinery, 2016), pp. 785–794.

22. M. López de Prado, Advances in Financial Machine Learning (Wiley, 2018).

23. S. M. Lundberg, S.-I. Lee, “A unified approach to interpreting model predictions” in Advances in Neural Information Processing Systems 30 (Curran Associates, 2017), pp. 4765–4774.

24. D. R. Cox, Regression models and life-tables. J. R. Stat. Soc. Ser. B Methodol. 34, 187–202 (1972).

25. P.-M. Lepretre, C. Foster, J.-P. Koralsztein, V. L. Billat, Heart rate deflection point as a strategy to defend stroke volume during incremental exercise. J. Appl. Physiol. 98, 1660–1665 (2005).

26. J. B. Crabbe, R. K. Dishman, Brain electrocortical activity during and after exercise: A quantitative synthesis. Psychophysiology 41, 563–574 (2004).

27. J. R. Manning, J. Jacobs, I. Fried, M. J. Kahana, Broadband shifts in local field potential power spectra are correlated with single-neuron spiking in humans. J. Neurosci. 29, 13613–13620 (2009).

28. N. Brake, F. Duc, A. Rokos, F. Arseneau, S. Shahiri, A. Khadra, G. Plourde, A neurophysiological basis for aperiodic EEG and the background spectral trend. Nat. Commun. 15, 1514 (2024).

29. S. D. Muthukumaraswamy, High-frequency brain activity and muscle artifacts in MEG/EEG: A review and recommendations. Front. Hum. Neurosci. 7, 138 (2013).

30. A. Wolff, N. Berberian, M. Golesorkhi, J. Gomez-Pilar, F. Zilio, G. Northoff, Intrinsic neural timescales: Temporal integration and segregation. Trends Cogn. Sci. 26, 159–173 (2022).

31. Y. Çatal, K. Keskin, A. Wolman, P. Klar, D. Smith, G. Northoff, Flexibility of intrinsic neural timescales during distinct behavioral states. Commun. Biol. 7, 1667 (2024).

32. D. Toker, I. Pappas, J. D. Lendner, J. Frohlich, D. M. Mateos, S. Muthukumaraswamy, R. Carhart-Harris, M. Paff, P. M. Vespa, M. M. Monti, F. T. Sommer, R. T. Knight, M. D’Esposito, Consciousness is supported by near-critical slow cortical electrodynamics. Proc. Natl. Acad. Sci. U.S.A. 119, e2024455119 (2022).

33. M. Schartner, A. Seth, Q. Noirhomme, M. Boly, M.-A. Bruno, S. Laureys, A. Barrett, Complexity of multi-dimensional spontaneous EEG decreases during propofol induced general anaesthesia. PLOS ONE 10, e0133532 (2015).

34. C. Avancini, C. Alameda, D. Sanabria, T. Bekinschtein, S. van Gaal, L. F. Ciria, Characterising the neural dynamics of physical exhaustion, Open Science Framework (2024); 10.17605/OSF.IO/TW5KV.

35. T. A. Bekinschtein, S. Dehaene, B. Rohaut, F. Tadel, L. Cohen, L. Naccache, Neural signature of the conscious processing of auditory regularities. Proc. Natl. Acad. Sci. U.S.A. 106, 1672–1677 (2009).

36. A. Delorme, S. Makeig, EEGLAB: An open source toolbox for analysis of single-trial EEG dynamics including independent component analysis. J. Neurosci. Methods 134, 9–21 (2004).

37. R. Oostenveld, P. Fries, E. Maris, J.-M. Schoffelen, FieldTrip: Open source software for advanced analysis of MEG, EEG, and invasive electrophysiological data. Comput. Intell. Neurosci. 2011, 156869 (2011).

38. L. S. Imperatori, M. Betta, L. Cecchetti, A. Canales-Johnson, E. Ricciardi, F. Siclari, P. Pietrini, S. Chennu, G. Bernardi, EEG functional connectivity metrics wPLI and wSMI account for distinct types of brain functional interactions. Sci. Rep. 9, 8894 (2019).

39. N. V. Chawla, K. W. Bowyer, L. O. Hall, W. P. Kegelmeyer, SMOTE: Synthetic minority over-sampling technique. J. Artif. Intell. Res. 16, 321–357 (2002).

40. C. Davidson-Pilon, lifelines: Survival analysis in Python. J. Open Source Softw. 4, 1317 (2019).

41. F. E. Harrell Jr, R. M. Califf, D. B. Pryor, K. L. Lee, R. A. Rosati, Evaluating the yield of medical tests. JAMA 247, 2543–2546 (1982).

