## Supplementary material for "Neural dynamics track the approach to physical exhaustion": Suplementary material: Supplementary material_bioRxiv_2026-11-12.pdf

**The PDF file includes:**

Supplementary Text
Figs. S1 to S17
Tables S1 to S13

#### 1. SUPPLEMENTARY TEXT

#### 1.1 Cardiorespiratory measures and physiological strain

We examined the association structure of the cardiorespiratory measures (Fig. 1C). All pairwise associations were positive and ranged from moderate to perfect ($\tau = 0.526$ to $1.000$). As expected, $VO_2$ and $VO_2$/kg were perfectly correlated ($\tau = 1.000$), and both showed strong associations with $VCO_2$ ($\tau = 0.935$) and HR ($\tau = 0.774$). $VCO_2$ was also strongly associated with HR ($\tau = 0.820$). RER showed comparatively weaker, although still moderate-to-strong, associations with the other physiological measures ($\tau = 0.526$ to $0.682$). Overall, this pattern indicated substantial redundancy among the cardiorespiratory variables.

To obtain a single index of physiological strain while avoiding multicollinearity, we performed PCA on the standardized cardiorespiratory variables. PC1 accounted for 87.5% of the total variance and showed uniformly positive loadings across all variables ($VO_2$ = 0.462, $VO_2$/kg = 0.462, $VCO_2$ = 0.478, RER = 0.327, HR = 0.488). The nearly uniform positive loadings indicate that PC1 represents the shared increase in cardiovascular and metabolic demand associated with progressive exercise intensity (Fig. 1C), rather than any single physiological variable and was therefore used as the physiological strain predictor in all subsequent analyses.

#### 1.2 Linear models

Piecewise linear models were used to characterize the relationship between physiological strain and neural activity across the increasing-effort and recovery phases (results summarised in Table S1 and Fig. 1D). For each neural measure, we compared a reduced model including participant identity and phase with a full model additionally including the physiological-strain component and its interaction with phase. Thus, the reported partial *F*-tests quantify whether physiological strain explained additional variance in the neural measure beyond participant and phase effects, while allowing the physiological–neural slope to differ between exercise and recovery. Statistical significance was assessed using family-wise max-statistic permutation p-values.

Physiological strain was significantly associated with all four oscillatory power bands. The strongest effect was observed for alpha power, ($F(2, 230) = 139.62$, $p < 0.001$, $\eta_p^2 = 0.548$), followed by delta power ($F(2, 230) = 124.09$, $p < 0.001$, $\eta_p^2 = 0.519$), and beta power ($F(2, 230) = 122.96$, $p < 0.001$, $\eta_p^2 = 0.517$). Theta power also showed a significant, although comparatively smaller, association ($F(2, 230) = 36.29$, $p < 0.001$, $\eta_p^2 = 0.240$). Inspection of the phase-specific estimates indicated that delta power was positively associated with physiological strain during both exercise and recovery, with a steeper estimated association during recovery (slopes = 0.245 and 0.457, respectively). In contrast, theta, alpha, and beta power were negatively associated with physiological strain in both phases, with more negative estimated slopes during recovery than during exercise (theta: -0.177 and -0.412; alpha: -0.188 and -0.600; beta: -0.223 and -0.676).

Both aperiodic parameters showed robust associations with physiological strain. The effect was stronger for the intercept ($F(2, 230) = 60.66$, $p < 0.001$, $\eta_p^2 = 0.345$), than for the slope ($F(2, 230) = 16.76$, $p = 0.001$, $\eta_p^2 = 0.127$). The intercept showed positive estimated associations with physiological strain during both exercise and recovery (slopes = 0.199 and 0.162, respectively). For the aperiodic slope parameter, the estimated association was close to zero during exercise but positive during recovery (slopes = 0.018 and 0.379 respectively).

All complexity measures showed significant associations with physiological strain. The largest effects were observed for KFD ($F(2, 230) = 50.57$, $p < 0.001$, $\eta_p^2 = 0.305$), and HFD ($F(2, 230) = 50.21$, $p < 0.001$, $\eta_p^2 = 0.304$). Significant effects were also found for FE ($F(2, 230) = 31.79$, $p < 0.001$, $\eta_p^2 = 0.217$), LZ ($F(2, 230) = 19.12$, $p < 0.001$, $\eta_p^2 = 0.143$), SE ($F(2, 230) = 16.75$, $p < 0.001$, $\eta_p^2 = 0.127$), AE ($F(2, 230) = 14.42$, $p = 0.001$, $\eta_p^2 = 0.111$), and KS ($F(2, 230) = 7.92$, $p = 0.026$, $\eta_p^2 = 0.064$). The phase-specific estimates revealed heterogeneous patterns. KFD and HFD were negatively associated with physiological strain during both phases, as was LZ, although its recovery slope was close to zero. AE and SE showed negative estimated associations during exercise but negligible associations during recovery. KS similarly changed from a small negative estimated association during exercise to a small positive association during recovery. In contrast, FE was positively associated with physiological strain during both phases, with a stronger estimated association during recovery.

Significant associations with physiological strain were also observed at all four wSMI time lags. The largest effect occurred at $\tau = 10$ ($F(2, 230) = 35.49$, $p < 0.001$, $\eta_p^2 = 0.236$) followed by $\tau = 4$ ($F(2, 230) = 28.10$, $p < 0.001$, $\eta_p^2 = 0.196$). Smaller but significant effects were found at $\tau = 40$ ($F(2, 230) = 8.12$, $p = 0.026$, $\eta_p^2 = .066$) and $\tau = 6$ ($F(2, 230) = 7.61$, $p = 0.037$, $\eta_p^2 = .062$). The estimated associations were negative at $\tau = 4$ during both exercise and recovery, positive at $\tau = 10$ and $\tau = 40$ during both phases, and close to zero during exercise but negative during recovery at $\tau = 6$.

### 1.3 Predictive models

The EEG features extracted from the near-exhaustion and far-from-exhaustion segments discriminated between the two classes with high accuracy across all participants and segment lengths tested. Discrimination was highest at the finest windows (AUROC of 0.992 ± 0.006 at 3 s and 0.989 ± 0.007 at 10 s) and decreased towards the coarsest windows (0.962 ± 0.021 at 30 s and 0.916 ± 0.056 at 60 s). When using 3-s segments, all 28 participants exceeded an AUROC of 0.95. Because the dataset was strongly class-imbalanced, the AUPRC provides a more stringent summary and followed the same pattern (0.969 ± 0.022 at 3 s to 0.853 ± 0.076 at 60 s). Threshold-dependent metrics, computed with the decision threshold fixed on the training folds, declined more steeply (F1 score 0.894 ± 0.034 at 3 s, 0.890 ± 0.046 at 10 s, and 0.740 ± 0.094 at 60 s). The fact that the threshold-dependent metrics (precision, recall and F1) fell more steeply than the ranking-based ones (AUROC and AUPRC) reflects the contraction of the near-exhaustion class at coarse windows: spanning the final five minutes, this class shrinks from roughly one hundred segments per participant at 3 s to about five at 60 s, destabilising both the minority-class oversampling and the per-fold estimation of precision and recall. The F1 decline was driven mainly by a loss of precision (0.849 ± 0.048 at 3 s to 0.683 ± 0.106 at 60 s) rather than of recall (0.951 ± 0.019 to 0.866 ± 0.076): with so few minority examples, the decision boundary became poorly calibrated and produced more false positives, even though the model continued to detect most true near-exhaustion segments. A second, signal-level factor compounds this for the EEG model: part of the discriminative neural information resides in fast spectral, complexity and connectivity dynamics that are averaged out over long windows, so longer segments lose signal as well as samples. Group-level metrics for all modalities and segment lengths are summarised in Table S2.

The cardiorespiratory features were also highly discriminative. They attained their highest accuracy at the finest window (AUROC 0.992 ± 0.012 and F1 0.931 ± 0.066 at 3 s), remained excellent up to 15 s (AUROC ≥ 0.985, F1 ≥ 0.904) and declined only modestly at the coarsest windows (AUROC 0.964 ± 0.021 and F1 0.862 ± 0.074 at 30 s; AUROC 0.928 ± 0.035 and F1 0.750 ± 0.088 at 60 s). The threshold-dependent metrics fell somewhat more than the ranking-based ones (precision 0.921 ± 0.074 at 3 s to 0.728 ± 0.098 at 60 s), reflecting the contraction of the near-exhaustion class at coarse windows. The modest size of this decline follows from the nature of the signal: the information that separates the two classes is carried by the sustained magnitude of the metabolic response (HR, $VO_2$ and RER rising towards exhaustion) a slow level that is faithfully represented even within a 3 s window and that keeps the two classes well separated, so short segments do not destabilise it and the decision boundary remains well calibrated. Its gradual decline at coarse windows is therefore attributable mainly to the statistical effect (the contraction of the near-exhaustion class and the associated reduction in the number of training segments) rather than to temporal averaging of the features themselves. The final minutes before exhaustion were therefore separable from the rest of the task in every participant and at every temporal scale examined, from three-second to sixty-second windows, indicating that the approach to the physical limit leaves a signature in cortical activity that is both individually reliable and robust to the temporal resolution at which it is measured.

#### 1.4 Feature importance analysis

To identify which neural features drove the classification, SHAP values were computed for the EEG model at the 10 s window on out-of-fold predictions under blocked cross-validation (gap 0 s) and aggregated as the per-participant normalised importance averaged across participants, with feature types ordered by their median (Fig. 2F). No single family dominated, but signal irregularity and band-limited power led the ranking. FE was by far the most influential feature type (33.8% of the total importance), followed by beta-band power (13.9%). HFD (6.3%), theta-band power (6.2%), alpha-band power (5.9%) and wSMI at the intermediate timescale ($\tau$ = 10, 5.8%) formed a second tier, and each remaining feature type contributed less than 5%. Grouped by family, complexity measures accounted for 51.9% of the total importance, band-limited power for 29.8%, connectivity for 10.6% and the aperiodic parameters for 7.7%. The apparent importance of connectivity was concentrated in a few participants with very large wSMI values rather than shared across the cohort: at $\tau$ = 10 the per-participant contribution ranged from near zero in most participants to over 60% in isolated cases, so that its cohort-median contribution was close to zero (Fig. 2F, individual points) while its mean was correspondingly inflated. FE, by contrast, ranked among the most important feature types at every segment length examined, as did beta-band power

(Fig. S16E–G). The importance profile was computed for the 3, 10, 15 and 30 s windows. The 60 s window was omitted because participants yielded too few segments at this resolution (median 25, maximum 39 per participant) for the blocked cross-validation scheme used to estimate the SHAP values, which requires at least 40 segments. Spatially, mean importance was highest over parieto-occipital, parietal and right temporal channels (PO7, PO3, T8, P6 and P5), although per-channel importance was heterogeneous across participants and its median was close to zero for all channels, so no single region dominated consistently. The classifier therefore relied chiefly on signal irregularity, captured most consistently by FE, together with band-limited power, rather than on cortical connectivity.

**1.5 Survival analysis**

Beyond the binary classification, a Cox PH model was used to characterise the continuous approach to exhaustion within each participant, providing both a discrimination metric (the C-index) and an estimate of temporal anticipation (the time remaining when the cumulative risk reached 80% of its final value, see Methods). The cardiorespiratory model offered a strong reference: at the 10 s window it reached a global C-index of 0.933 ± 0.023 and reached the 80% level a median of 40 s before cessation (IQR 23–94 s; Fig. 3G; per-window results in Table S5). Consistent with the established physiological literature, cardiorespiratory dynamics therefore tracked the progression towards the physical limit with high fidelity.

The EEG-only model reached a global C-index of 0.897 ± 0.058 at the same window and reached the 80% level a median of 60 s before cessation (IQR 33–73 s; per-window results in Table S4). The neural signal alone thus carried information about the proximity to exhaustion of a magnitude comparable to that of the cardiorespiratory model, indicating that the approach to the physical limit is reflected in cortical activity and not only in peripheral physiological responses. Because the early portion of the protocol contributes little to the prediction of imminent exhaustion (an experienced athlete does not approach cessation during the initial minutes) model performance was additionally examined over the final phase of the task. When performance in this final phase was quantified by whether segments in the terminal 25% of the task were ranked above earlier segments, both modalities retained near-perfect discrimination (late-versus-rest C-index 0.959 ± 0.090 for EEG and 0.981 ± 0.026 for the cardiorespiratory model), consistent with the time-resolved curves in Fig. 3I. As a more demanding stress test, concordance was further restricted to pairs of segments both lying within the terminal 25% (that is, the ordering of physiologically near-identical, adjacent states immediately before exhaustion).

Under this deliberately hard criterion, the two models dropped to more modest levels (final-phase C-index 0.736 ± 0.105 for EEG and 0.701 ± 0.200 for the cardiorespiratory model), reflecting the intrinsic difficulty of resolving fine temporal order within a near-exhaustion plateau rather than a failure of the models. In this regime the cardiorespiratory estimate also showed markedly greater inter-participant variability than the EEG estimate. The temporal performance curves (Fig. 3I) illustrate the same pattern: the three models tracked the task comparably for most of its duration, whereas in the segments immediately preceding exhaustion the cardiorespiratory curve declined and its confidence band widened, while the EEG curve remained stable. These within-participant estimates were not materially inflated by fitting and evaluating on the same data: when the Cox model was fitted on contiguous training blocks and evaluated on held-out blocks with a 10 s gap, the EEG C-index was 0.871 ± 0.069 (final quarter 0.664 ± 0.157), against 0.897 in sample (Table S10). By contrast, a model trained on the other participants transferred less well to EEG (LOPO C-index 0.781 ± 0.081) than to cardiorespiratory signals (0.916 ± 0.059), and the group-derived EEG risk trajectory lost its characteristic late rise (Fig. S17B and C, Table S12), indicating that the cortical signature of approaching exhaustion is largely participant-specific.

The combined model, which joined EEG and cardiorespiratory features by early fusion, retained the highest final-phase discrimination (0.800 ± 0.099) and reached the 80% level closest to cessation (median 27 s, IQR 17–59 s; EEG 60 s, cardiorespiratory 40 s; Friedman $\chi^2(2) = 19.8$, $p = 5.1 \times 10^{-5}$). Its warning came later than the cardiorespiratory one in 26 of 28 participants and later than the EEG one in 21 of 28 (Holm-corrected Wilcoxon, both $p \leq 0.012$), whereas the two unimodal models did not differ ($p = 0.90$). Fusion also reduced the dispersion of anticipation across participants (sign-flip permutation on absolute deviations from the median, combined versus cardiorespiratory model, $p = 0.003$), and the result held at the 70% and 90% levels (Table S11). The results for the 10 s window are summarised in Table S3; the remaining segment lengths and an asynchronous fusion variant followed the same pattern (Tables S4–S7, Fig. S16A–D and Fig. S17A).

To confirm that this advantage of the combined model reflected genuine information rather than the larger dimensionality of the fused model, the permutation test described in Methods was applied at the 10 s window (10,000 permutations per participant). Adding the EEG features to the cardiorespiratory model improved discrimination beyond what dimensionality-matched, temporally permuted EEG achieved, both over all segment pairs (gain +0.016, significant in 25 of 28 participants; Stouffer $Z = 17.1$, $p < 3.5 \cdot 10^{-66}$) and within the final 25% of the task (gain +0.099, significant in 19 of 28 participants; Stouffer $Z = 11.7$, $p < 9.7 \cdot 10^{-32}$). The neural contribution was statistically robust across the cohort, indicating that the EEG carried information complementary to the cardiorespiratory signal rather than an artefact of model dimensionality. As expected, over the whole exercise the cardiorespiratory variables remained the dominant source of discrimination: adding them to the EEG model produced a larger gain than the reverse (over all pairs, +0.052 versus +0.016). Within the final quarter of the task, however, this ordering inverted: adding temporally aligned EEG to the cardiorespiratory model yielded a larger gain than the reverse (+0.099 versus +0.064), indicating that near the physical limit the neural signal contributes discrimination beyond, rather than merely below, the cardiorespiratory one. The neural contribution over all segment pairs was stable across segment lengths (+0.016 at 3 s, significant in 26 of 28 participants; +0.016 at 10 s, 25/28; +0.017 at 15 s, 25/28; +0.015 at 30 s, 19/28; +0.012 at 60 s, 12/28) and significant at the cohort level at every length (Stouffer $Z = 6.0$–$17.6$). Within the final 25% the gain was five to ten times larger (+0.080 to +0.116), but the number of participants in whom it exceeded the null fell from 23 of 28 at 3 s to 7 of 28 at 60 s. At 60 s the cohort-level test remained significant, although weakly (Stouffer $Z = 2.3$, $p = 0.0095$), because only about six segments fall within the final quarter at this resolution and few concordant pairs remain (Table S8). The complementary neural information is therefore best resolved at fine segment lengths.

The per-participant trajectories (Figs. S1–S15) showed why fusion produced the most consistent warnings. Each unimodal model produced early warnings in a few participants whose cumulative risk rose gradually rather than sharply, typically those in whom that model discriminated poorly (lowest per-participant C-index 0.63 for EEG and 0.88 for the cardiorespiratory model). Under the 80% criterion, the earliest warning came 4.7 min before cessation for EEG and 4.1 min for the cardiorespiratory model, and more than two minutes before cessation in 2 and 5 participants, respectively. The two modalities did not fail in the same participants, and fusion recovered a timely warning where one modality failed: in the participant with the earliest cardiorespiratory warning

(4.1 min), the combined model signalled at 1.5 min (EEG alone: 1.4 min); in the participant with the earliest EEG warning (4.7 min, C-index 0.63), the combined model signalled at 19 s, matching the cardiorespiratory component. The combined model discriminated well in every participant (minimum C-index 0.92) and warned within the final two minutes in 26 of 28 participants.

### 2. SUPPLEMENTARY FIGURES S1–S17

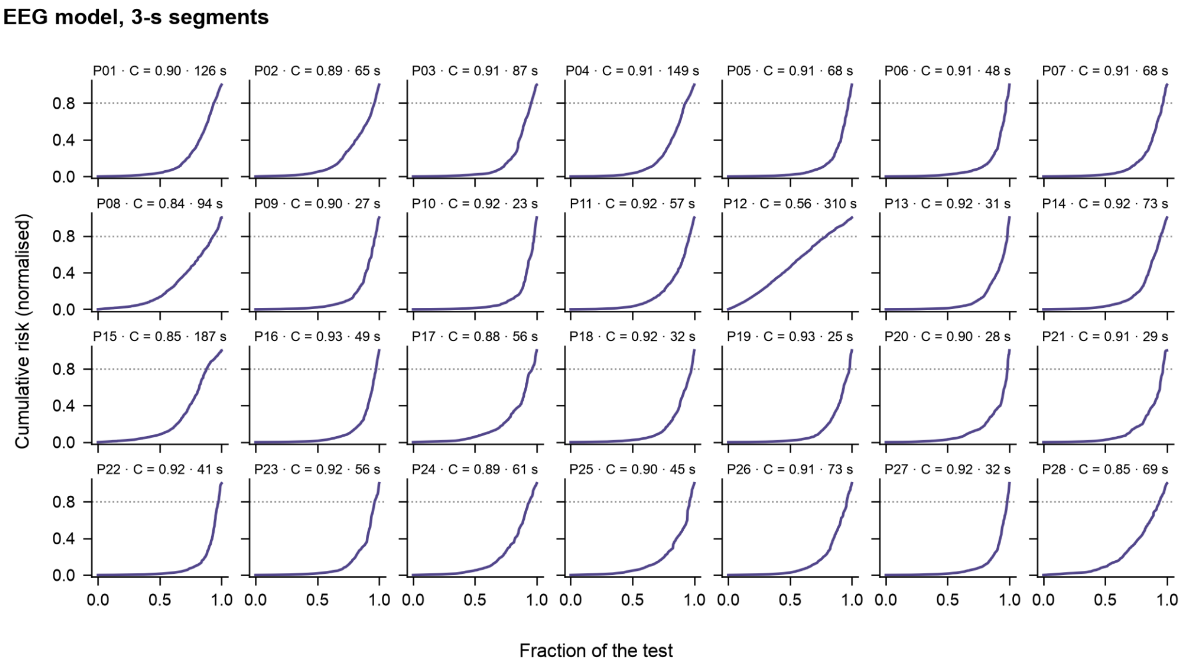


**Figure S1.** Per-participant cumulative-risk trajectories of the within-participant EEG Cox model (3 s window, n = 28). Each panel corresponds to one participant; the curve is the normalised cumulative risk across the exercise timeline (0 = start, 1 = exhaustion). The dotted horizontal line marks 80% of the final value and the dashed vertical line the point at which it was reached. Panel titles give the participant, the C-index and the anticipation (s).

**EEG model, 10-s segments**

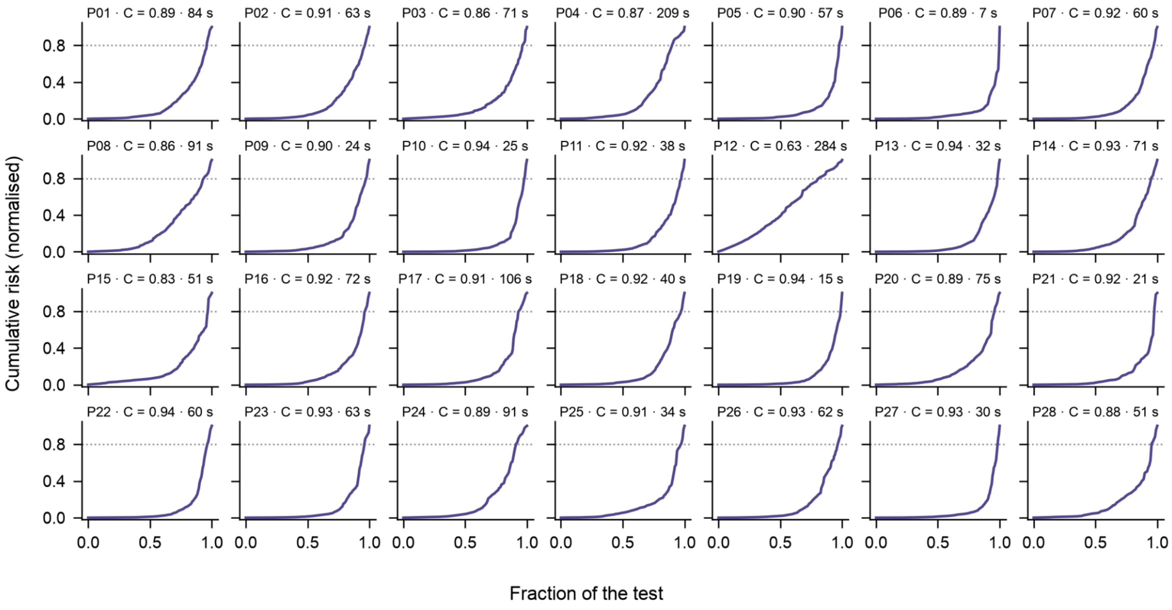


**Figure S2.** Per-participant cumulative-risk trajectories of the within-participant EEG Cox model (10 s window, n = 28). Each panel corresponds to one participant; the curve is the normalised cumulative risk across the exercise timeline (0 = start, 1 = exhaustion). The dotted horizontal line marks 80% of the final value and the dashed vertical line the point at which it was reached. Panel titles give the participant, the C-index and the anticipation (s).

**EEG model, 15-s segments**

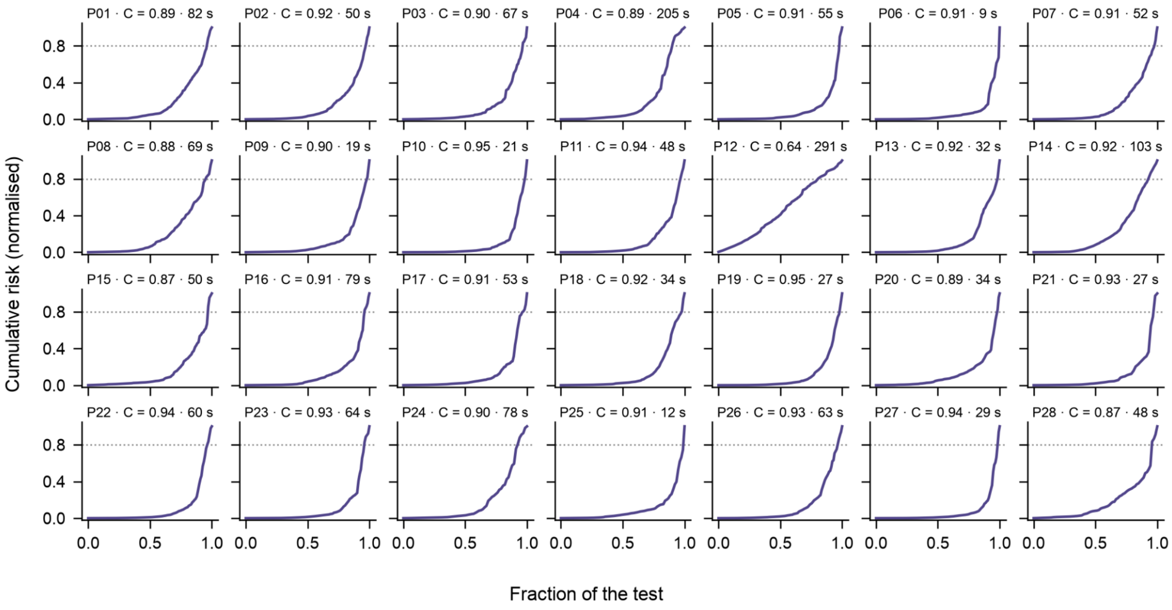


**Figure S3.** Per-participant cumulative-risk trajectories of the within-participant EEG Cox model (15 s window, n = 28). Each panel corresponds to one participant; the curve is the normalised cumulative risk across the exercise timeline (0 = start, 1 = exhaustion). The dotted horizontal line marks 80% of the final value and the dashed vertical line the point at which it was reached. Panel titles give the participant, the C-index and the anticipation (s).

### EEG model, 30-s segments

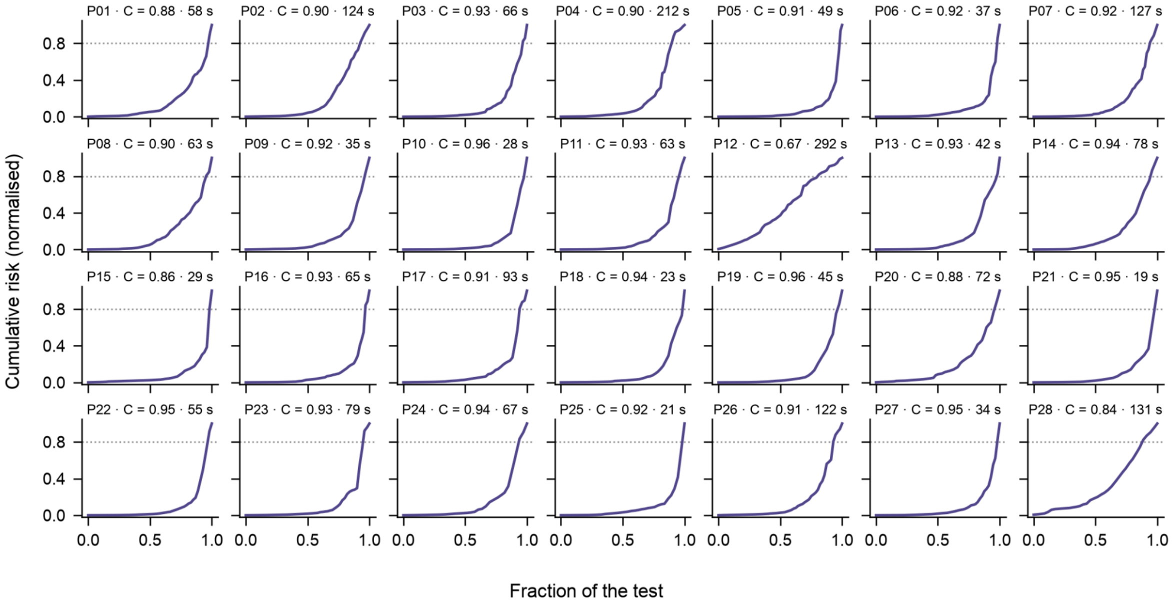


**Figure S4.** Per-participant cumulative-risk trajectories of the within-participant EEG Cox model (30 s window, n = 28). Each panel corresponds to one participant; the curve is the normalised cumulative risk across the exercise timeline (0 = start, 1 = exhaustion). The dotted horizontal line marks 80% of the final value and the dashed vertical line the point at which it was reached. Panel titles give the participant, the C-index and the anticipation (s).

### EEG model, 60-s segments

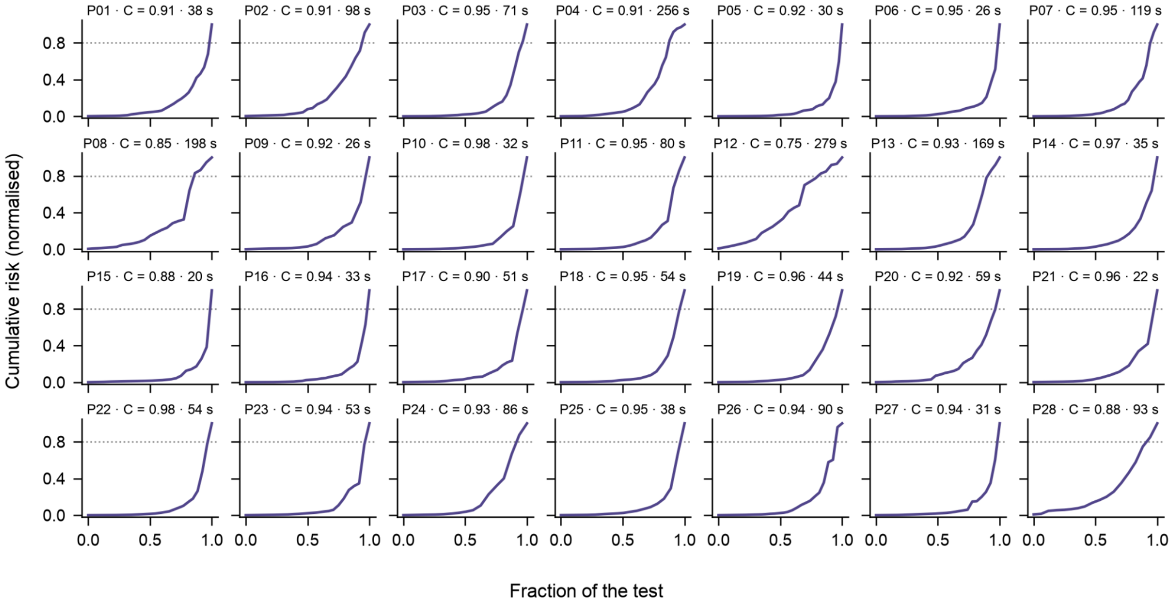


**Figure S5.** Per-participant cumulative-risk trajectories of the within-participant EEG Cox model (60 s window, n = 28). Each panel corresponds to one participant; the curve is the normalised cumulative risk across the exercise timeline (0 = start, 1 = exhaustion). The dotted horizontal line marks 80% of the final value and the dashed vertical line the point at which it was reached. Panel titles give the participant, the C-index and the anticipation (s).

#### Cardiorespiratory model, 3-s segments

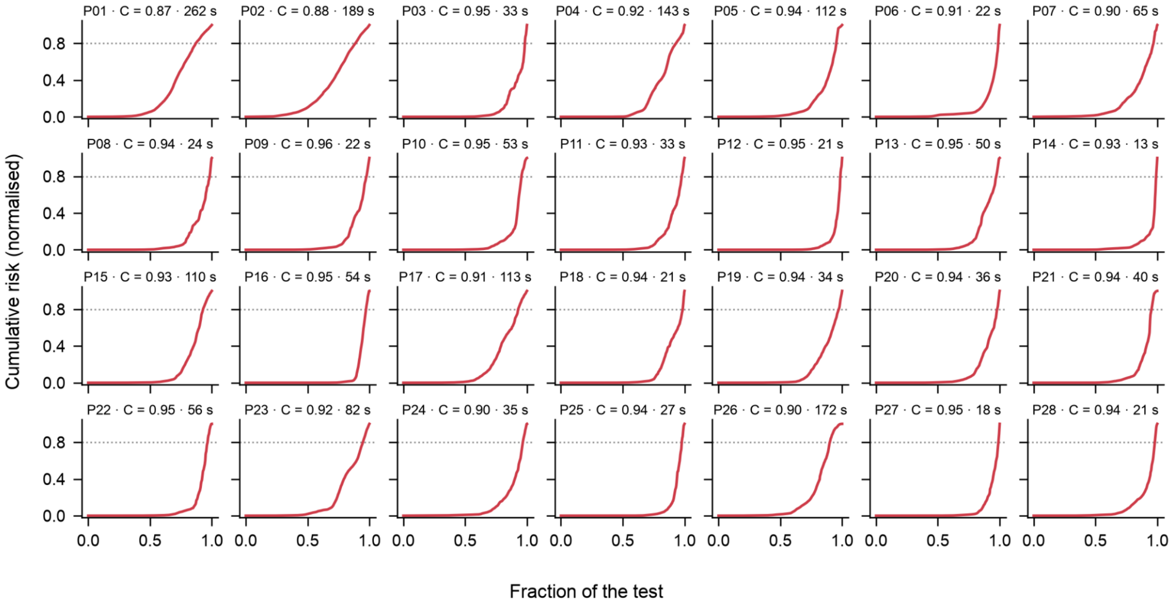


**Figure S6.** Per-participant cumulative-risk trajectories of the within-participant cardiorespiratory Cox model (3 s window, n = 28). Each panel corresponds to one participant; the curve is the normalised cumulative risk across the exercise timeline (0 = start, 1 = exhaustion). The dotted horizontal line marks 80% of the final value and the dashed vertical line the point at which it was reached. Panel titles give the participant, the C-index and the anticipation (s).

#### Cardiorespiratory model, 10-s segments

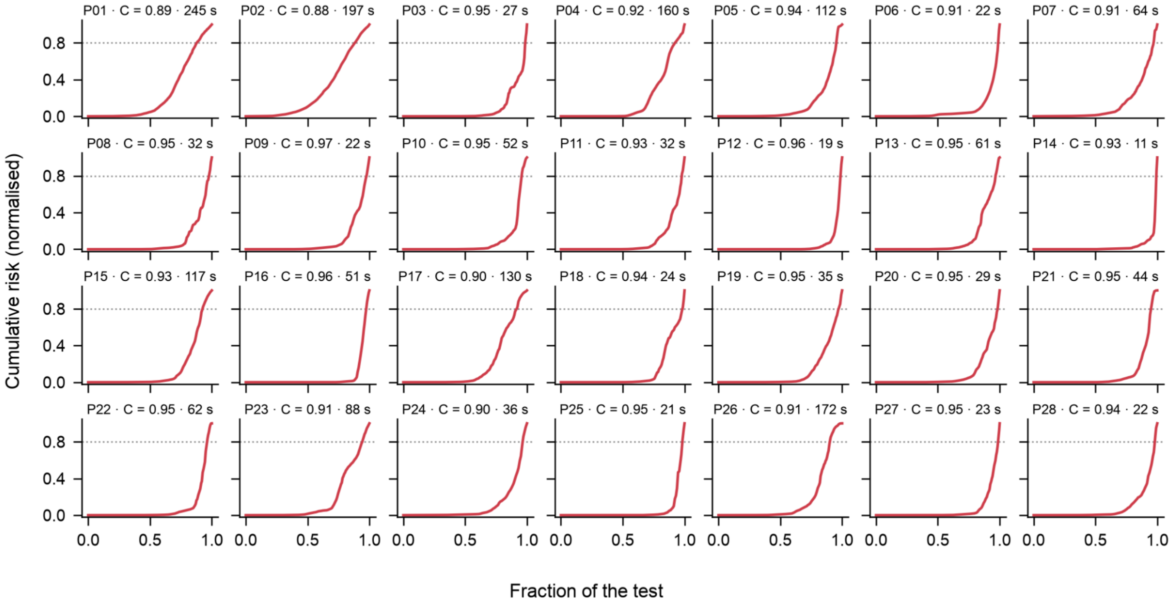


**Figure S7.** Per-participant cumulative-risk trajectories of the within-participant cardiorespiratory Cox model (10 s window, n = 28). Each panel corresponds to one participant; the curve is the normalised cumulative risk across the exercise timeline (0 = start, 1 = exhaustion). The dotted horizontal line marks 80% of the final value and the dashed vertical line the point at which it was reached. Panel titles give the participant, the C-index and the anticipation (s).

**Cardiorespiratory model, 15-s segments**

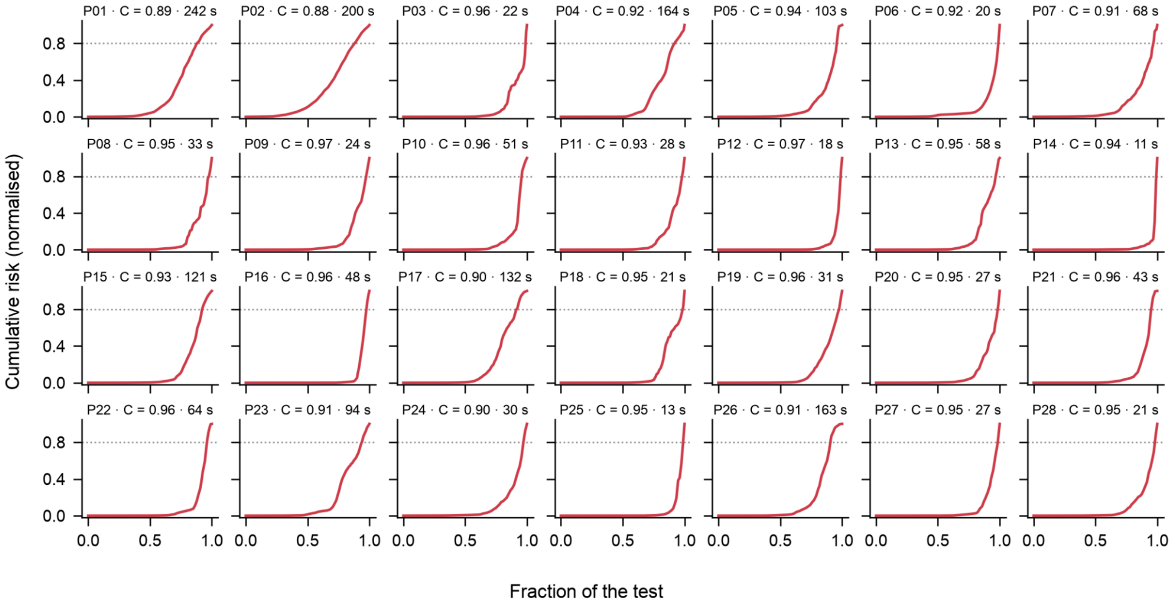


**Figure S8**. Per-participant cumulative-risk trajectories of the within-participant cardiorespiratory Cox model (15 s window, n = 28). Each panel corresponds to one participant; the curve is the normalised cumulative risk across the exercise timeline (0 = start, 1 = exhaustion). The dotted horizontal line marks 80% of the final value and the dashed vertical line the point at which it was reached. Panel titles give the participant, the C-index and the anticipation (s).

**Cardiorespiratory model, 30-s segments**

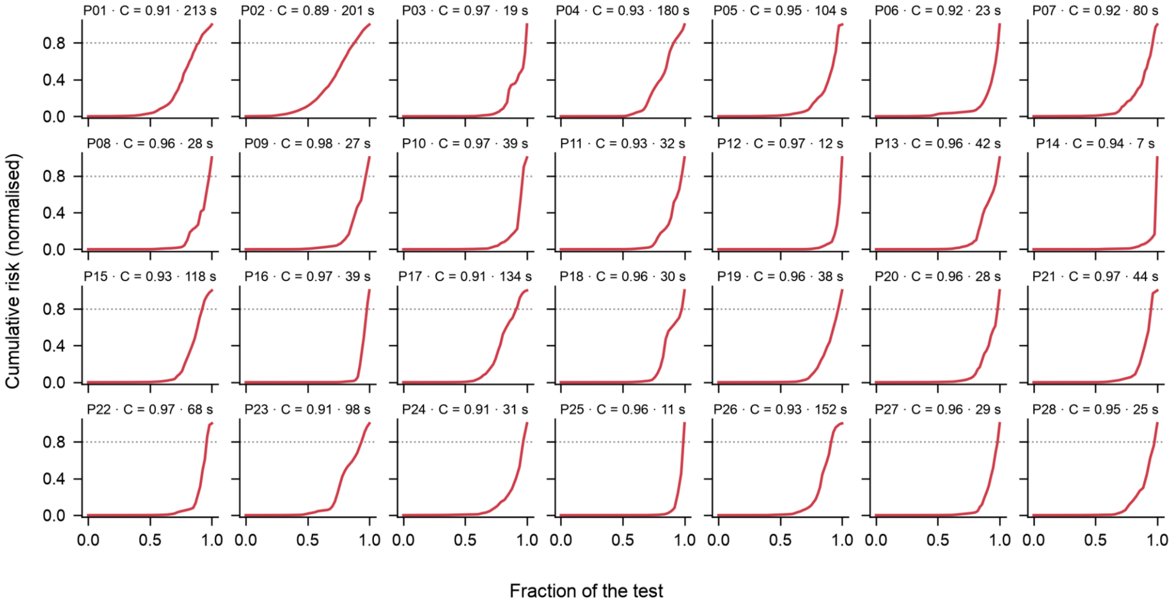


**Figure S9**. Per-participant cumulative-risk trajectories of the within-participant cardiorespiratory Cox model (30 s window, n = 28). Each panel corresponds to one participant; the curve is the normalised cumulative risk across the exercise timeline (0 = start, 1 = exhaustion). The dotted horizontal line marks 80% of the final value and the dashed vertical line the point at which it was reached. Panel titles give the participant, the C-index and the anticipation (s).

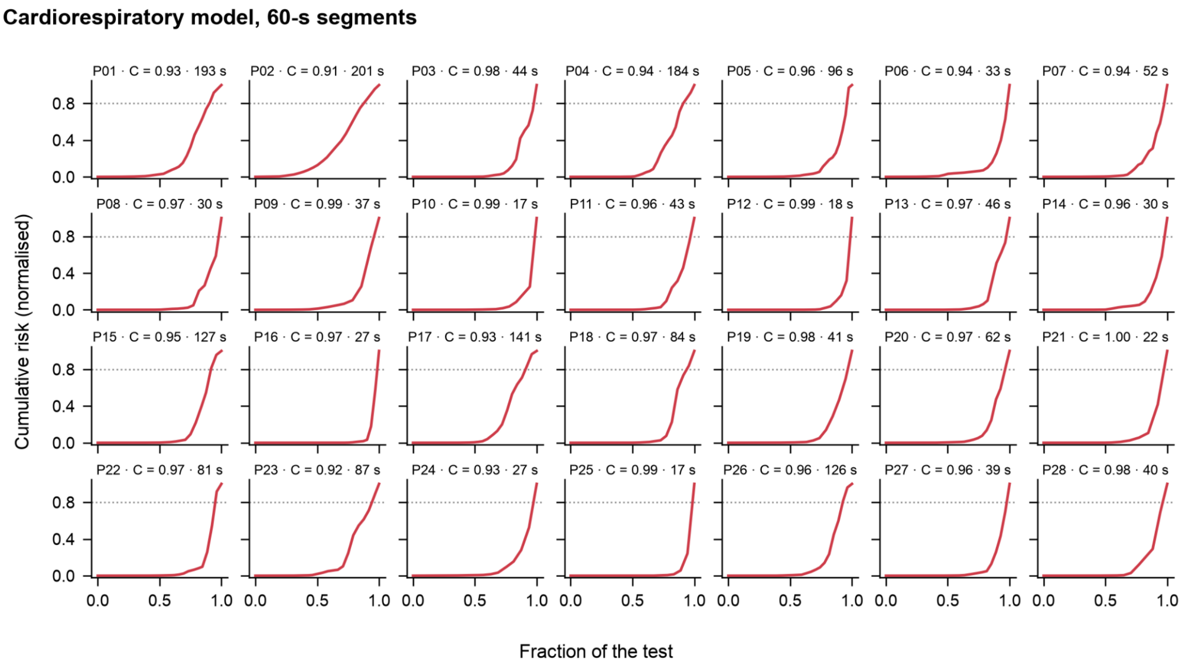


**Figure S10**. Per-participant cumulative-risk trajectories of the within-participant cardiorespiratory Cox model (60 s window, n = 28). Each panel corresponds to one participant; the curve is the normalised cumulative risk across the exercise timeline (0 = start, 1 = exhaustion). The dotted horizontal line marks 80% of the final value and the dashed vertical line the point at which it was reached. Panel titles give the participant, the C-index and the anticipation (s).

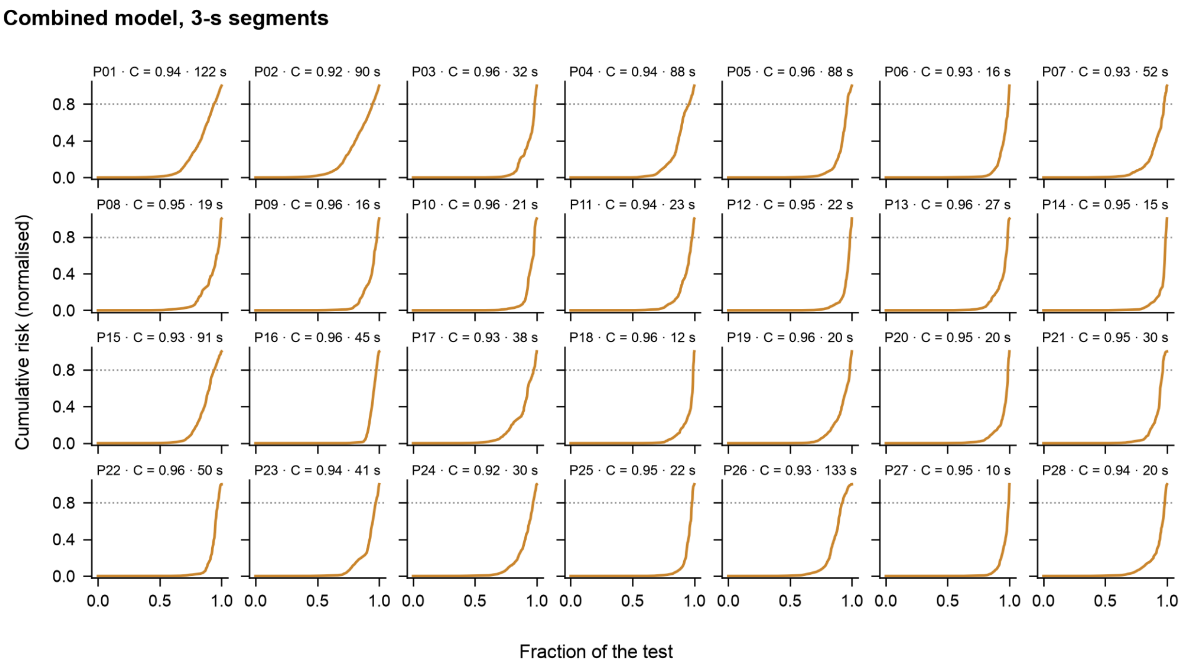


**Figure S11**. Per-participant cumulative-risk trajectories of the within-participant combined (EEG + cardiorespiratory, early fusion) Cox model (3 s window, n = 28). Each panel corresponds to one participant; the curve is the normalised cumulative risk across the exercise timeline (0 = start, 1 = exhaustion). The dotted horizontal line marks 80% of the final value and the dashed vertical line the point at which it was reached. Panel titles give the participant, the C-index and the anticipation (s).

**Combined model, 10-s segments**

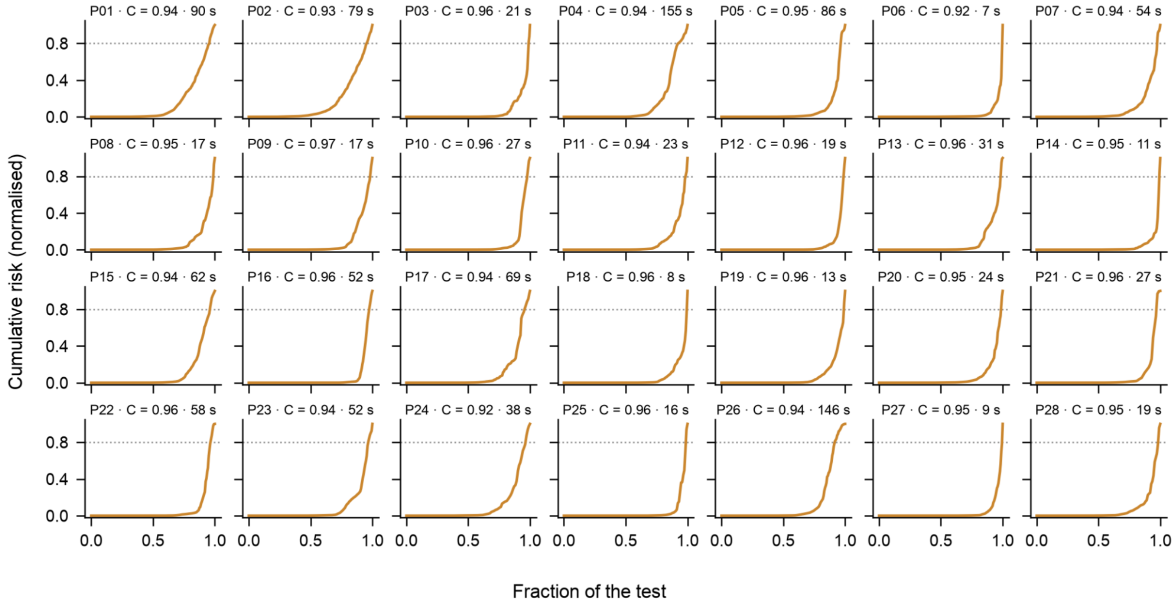


**Figure S12**. Per-participant cumulative-risk trajectories of the within-participant combined (EEG + cardiorespiratory, early fusion) Cox model (10 s window, n = 28). Each panel corresponds to one participant; the curve is the normalised cumulative risk across the exercise timeline (0 = start, 1 = exhaustion). The dotted horizontal line marks 80% of the final value and the dashed vertical line the point at which it was reached. Panel titles give the participant, the C-index and the anticipation (s).

**Combined model, 15-s segments**

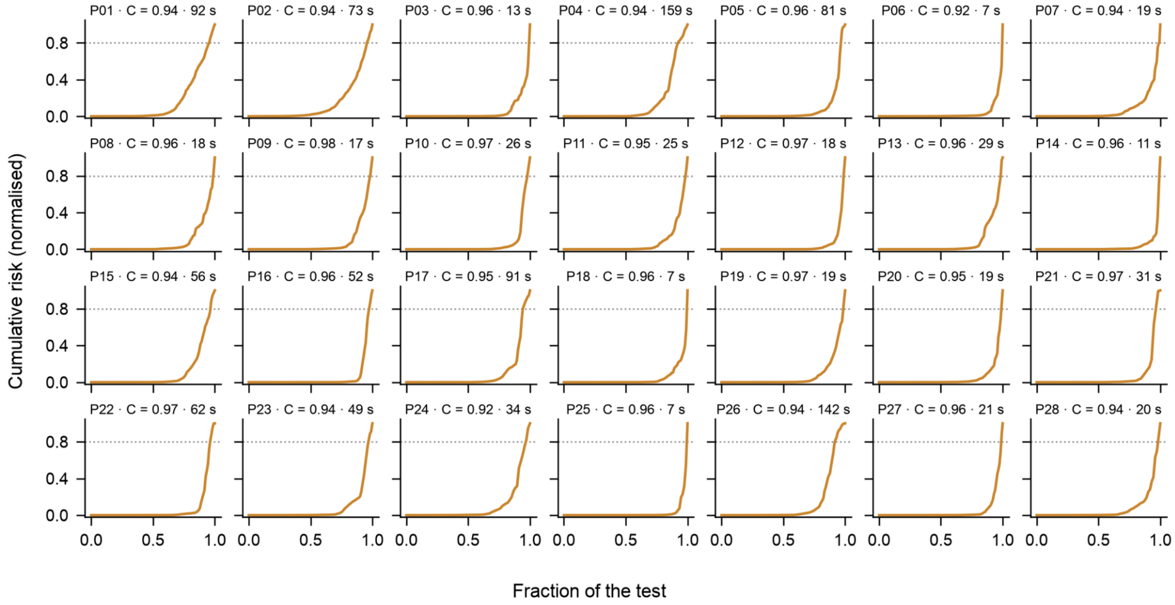


**Figure S13.** Per-participant cumulative-risk trajectories of the within-participant combined (EEG + cardiorespiratory, early fusion) Cox model (15 s window, n = 28). Each panel corresponds to one participant; the curve is the normalised cumulative risk across the exercise timeline (0 = start, 1 = exhaustion). The dotted horizontal line marks 80% of the final value and the dashed vertical line the point at which it was reached. Panel titles give the participant, the C-index and the anticipation (s).

**Combined model, 30-s segments**

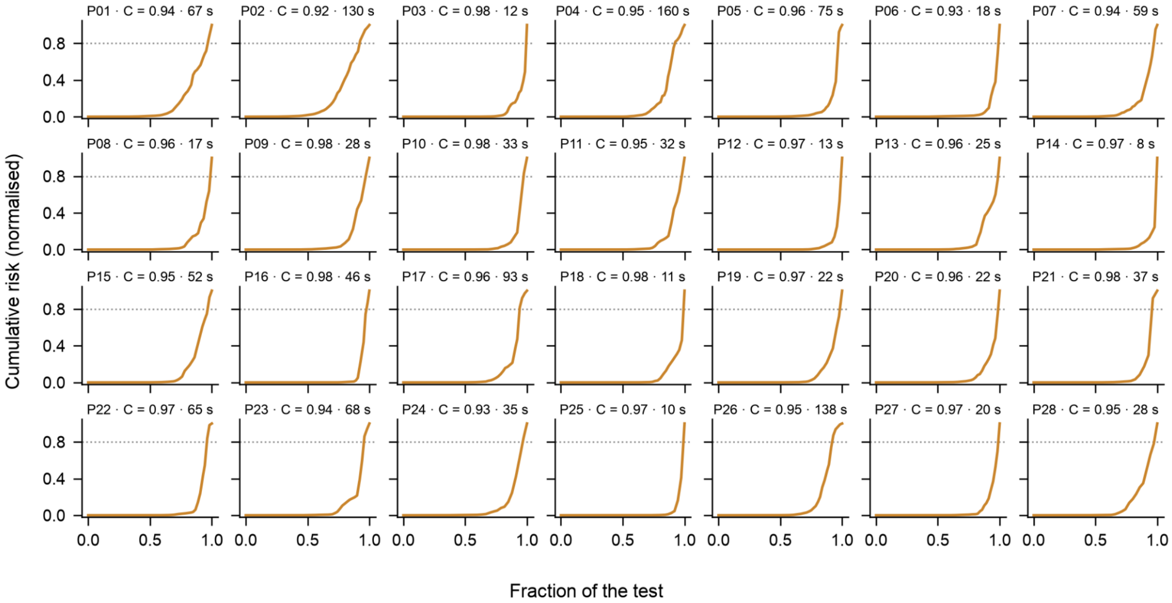


**Figure S14.** Per-participant cumulative-risk trajectories of the within-participant combined (EEG + cardiorespiratory, early fusion) Cox model (30 s window, n = 28). Each panel corresponds to one participant; the curve is the normalised cumulative risk across the exercise timeline (0 = start, 1 = exhaustion). The dotted horizontal line marks 80% of the final value and the dashed vertical line the point at which it was reached. Panel titles give the participant, the C-index and the anticipation (s).

**Combined model, 60-s segments**

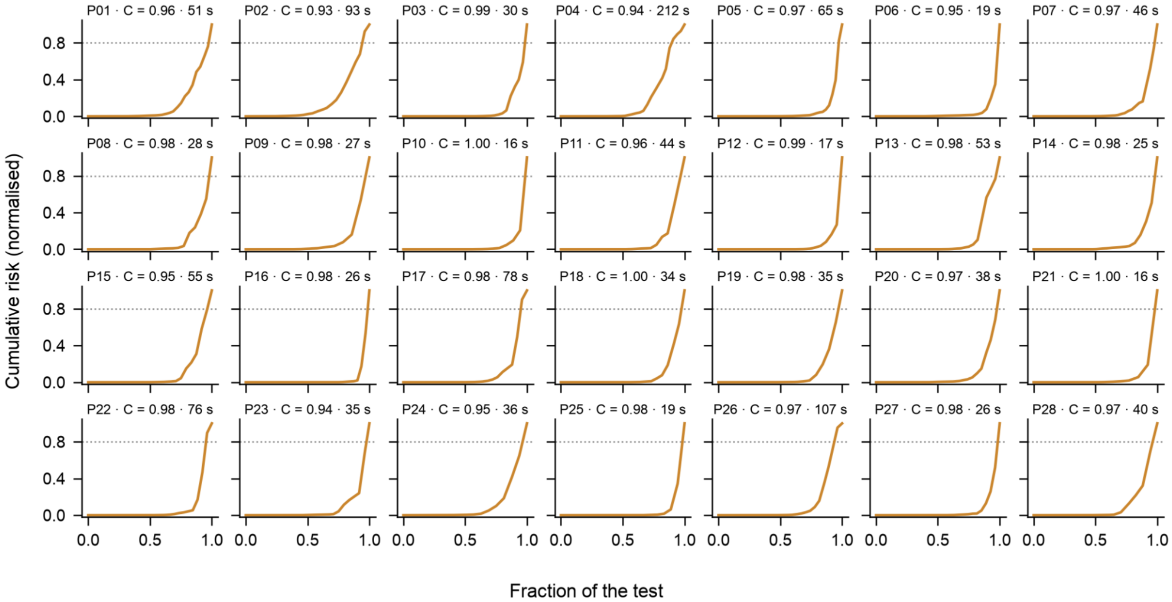


**Figure S15.** Per-participant cumulative-risk trajectories of the within-participant combined (EEG + cardiorespiratory, early fusion) Cox model (60 s window, n = 28). Each panel corresponds to one participant; the curve is the normalised cumulative risk across the exercise timeline (0 = start, 1 = exhaustion). The dotted horizontal line marks 80% of the final value and the dashed vertical line the point at which it was reached. Panel titles give the participant, the C-index and the anticipation (s).

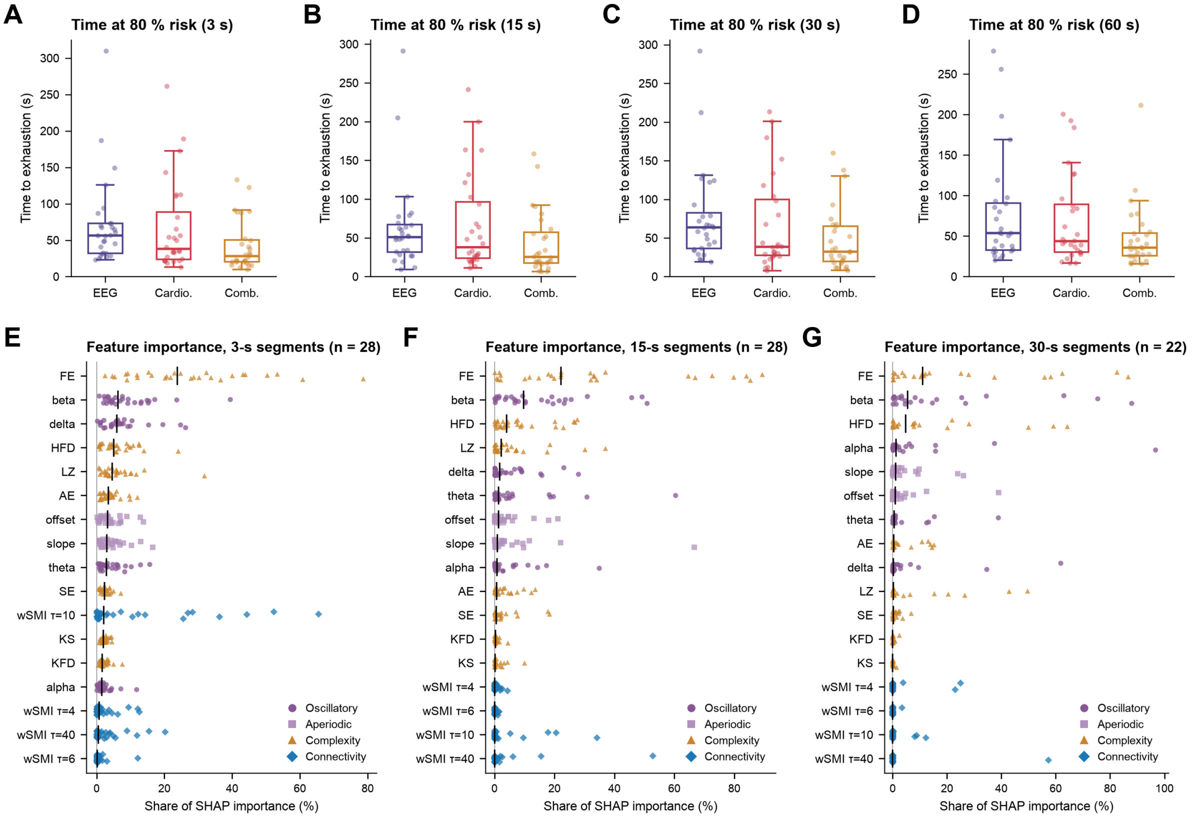


**Figure S16.** Anticipation and feature importance at the remaining segment lengths. (A–D) Anticipation (time remaining when the cumulative risk reached 80% of its final value) for the EEG, cardiorespiratory and combined within-participant Cox models at the 3, 15, 30 and 60 s windows (n = 28), presented as in Fig. 3G. Points are participants and boxes show the median and interquartile range. (E–G) Feature-type importance of the EEG classifier at the 3, 15 and 30 s windows (n = 28, 28 and 22), presented as in Fig. 2F. Each point is one participant's share of SHAP importance for a feature type (mean absolute SHAP value expressed as a percentage of the sum across features within the participant). Black lines mark the median across participants, feature types are ordered by their median share and colours indicate the feature family. FE carried the largest share at every window (mean 27.3%, 32.5% and 23.3% at 3, 15 and 30 s). It was followed by wSMI at τ = 10 at 3 s (12.1%, carried by a few participants with very large values) and by beta-band power at 15 and 30 s (14.9% and 17.7%). The 60 s window is not shown in E–G because it yielded too few segments for the blocked cross-validation.

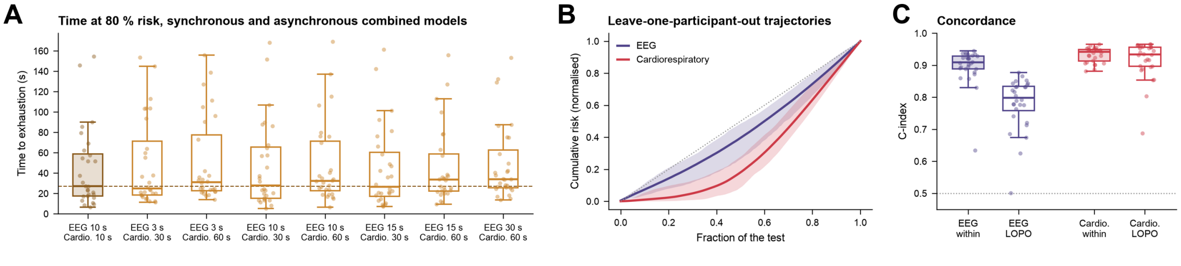


**Figure S17.** Combined models with asynchronous windows and between-participant transfer (n = 28). (A) Anticipation (80% criterion) for the synchronous combined model (10 s) and the seven asynchronous combinations (EEG window 3–30 s paired with a cardiorespiratory window of 30 or 60 s), presented as in Fig. 3G. Points are participants and boxes show the median and interquartile range. Median anticipation stayed between 25 and 34 s and 25–26 of 28 participants were warned within the final two minutes in every pairing (Table S7), so the combined warning does not depend on the two modalities sharing a temporal resolution. (B) Cumulative-risk trajectories of held-out participants for EEG and cardiorespiratory Cox models (10 s window) trained on the remaining 27 participants (leave-one-participant-out), expressed as a function of the fraction of the test completed. Lines show the mean across held-out participants and shaded bands the 2.5th–97.5th percentile range. The dotted line is the identity, corresponding to risk accumulating uniformly over the test. (C) Per-participant C-index of the within-participant models (filled boxes) and under leave-one-participant-out transfer (open boxes). Points are participants and the dotted line marks chance level. The cardiorespiratory model transferred almost intact (0.933 → 0.916), whereas EEG concordance dropped (0.897 → 0.781) and the group-derived EEG trajectory lost the late rise characteristic of the participant-specific models.

### 3. SUPPLEMENTARY TABLES S1–S13

| Family | Measure | F | p-value | ΔR² | $\eta_p^2$ | Interaction β | Slope exercise | Slope recovery |
|---|---|---|---|---|---|---|---|---|
| Oscillatory | delta | 124.093 | 0.000 | 0.261 | 0.519 | 0.212 | 0.245 | 0.457 |
| Oscillatory | theta | 36.292 | 0.000 | 0.159 | 0.240 | -0.235 | -0.177 | -0.412 |
| Oscillatory | alpha | 139.620 | 0.000 | 0.241 | 0.548 | -0.412 | -0.188 | -0.600 |
| Oscillatory | beta | 122.957 | 0.000 | 0.321 | 0.517 | -0.453 | -0.223 | -0.676 |
| Aperiodic | slope | 16.757 | 0.000 | 0.054 | 0.127 | 0.361 | 0.018 | 0.379 |
| Aperiodic | intercept | 60.661 | 0.000 | 0.132 | 0.345 | -0.038 | 0.199 | 0.162 |
| Complexity | AE | 14.419 | 0.001 | 0.074 | 0.111 | 0.168 | -0.155 | 0.013 |
| Complexity | FE | 31.789 | 0.000 | 0.109 | 0.217 | 0.417 | 0.079 | 0.495 |
| Complexity | KS | 7.918 | 0.026 | 0.043 | 0.064 | 0.257 | -0.107 | 0.149 |
| Complexity | SE | 16.746 | 0.000 | 0.085 | 0.127 | 0.176 | -0.166 | 0.010 |
| Complexity | HFD | 50.210 | 0.000 | 0.199 | 0.304 | 0.026 | -0.244 | -0.217 |
| Complexity | KFD | 50.568 | 0.000 | 0.222 | 0.305 | -0.051 | -0.248 | -0.299 |
| Complexity | LZ | 19.122 | 0.000 | 0.094 | 0.143 | 0.158 | -0.175 | -0.017 |
| wSMI | τ 4 | 28.100 | 0.000 | 0.103 | 0.196 | -0.121 | -0.156 | -0.278 |
| wSMI | τ 6 | 7.614 | 0.037 | 0.038 | 0.062 | -0.204 | -0.063 | -0.267 |
| wSMI | τ 10 | 35.490 | 0.000 | 0.140 | 0.236 | 0.022 | 0.199 | 0.221 |
| wSMI | τ 40 | 8.123 | 0.026 | 0.037 | 0.066 | 0.053 | 0.097 | 0.150 |

**Table S1. Cardiorespiratory–neural coupling across increasing exertion and recovery.** Results of the piecewise linear models with physiological strain, indexed by the first principal component (PC1) of the cardiorespiratory measures, as predictor and to each neural outcome as predicted variable. The *F* statistic compares the full model (including physiological strain and its interaction with protocol phase) with a reduced model including participant and phase only; all tests had 2 and 230 degrees of freedom. Reported *p* values are family-wise error–corrected within each neural family using a maximum-statistic permutation procedure with 10,000 permutations. $\Delta R^2$ indicates the additional variance explained by the physiological predictors, and $\eta_p^2$ quantifies their effect size. Interaction β is the coefficient for the physiological-strain × phase interaction. Exercise and recovery slopes are estimated associations between physiological strain and the neural outcome within each phase. For wSMI, τ indicates the temporal separation in samples.

| Segment | AUROC | AUPRC | Precision | Recall | F1 |
|---|---|---|---|---|---|
| **EEG** | | | | | |
| 3 s | 0.992 ± 0.006 | 0.969 ± 0.022 | 0.849 ± 0.048 | 0.951 ± 0.019 | 0.894 ± 0.034 |
| 10 s | 0.989 ± 0.007 | 0.966 ± 0.020 | 0.864 ± 0.060 | 0.936 ± 0.028 | 0.890 ± 0.046 |
| 15 s | 0.984 ± 0.009 | 0.959 ± 0.021 | 0.856 ± 0.049 | 0.928 ± 0.034 | 0.877 ± 0.042 |
| 30 s | 0.962 ± 0.021 | 0.916 ± 0.033 | 0.815 ± 0.056 | 0.901 ± 0.051 | 0.829 ± 0.049 |
| 60 s | 0.916 ± 0.056 | 0.853 ± 0.076 | 0.683 ± 0.106 | 0.866 ± 0.076 | 0.740 ± 0.094 |
| **Cardiorespiratory** | | | | | |
| 3 s | 0.992 ± 0.012 | 0.966 ± 0.055 | 0.921 ± 0.074 | 0.945 ± 0.056 | 0.931 ± 0.066 |
| 10 s | 0.987 ± 0.013 | 0.955 ± 0.050 | 0.915 ± 0.073 | 0.923 ± 0.062 | 0.911 ± 0.071 |
| 15 s | 0.985 ± 0.011 | 0.954 ± 0.042 | 0.912 ± 0.074 | 0.920 ± 0.048 | 0.904 ± 0.063 |
| 30 s | 0.964 ± 0.021 | 0.921 ± 0.040 | 0.883 ± 0.084 | 0.883 ± 0.059 | 0.862 ± 0.074 |
| 60 s | 0.928 ± 0.035 | 0.867 ± 0.065 | 0.728 ± 0.098 | 0.799 ± 0.072 | 0.750 ± 0.088 |

**Table S2. Binary classification performance of the EEG (64 channels, 836 features) and cardiorespiratory (5 features) models across segment lengths and modalities.** Values represent mean ± SD across participants (n = 28). AUROC: area under the receiver operating characteristic curve; AUPRC: area under the precision–recall curve. All 28 participants contributed valid estimates at every window length.

| Model | Global C-index | Final-phase C-index (last 25%) | Anticipation $t_{80}$ (s) | ≤ 2min |
|---|---|---|---|---|
| EEG | 0.897 ± 0.058 | 0.736 ± 0.105 | 60 [33-73] | 26/28 |
| Cardiorespiratory | 0.933 ± 0.023 | 0.701 ± 0.200 | 40 [23-94] | 23/28 |
| Combined | 0.949 ± 0.014 | 0.800 ± 0.099 | 27 [17-59] | 26/28 |

**Table S3. Within-participant Cox survival-analysis performance for the three modalities at the 10 s segment window** (n = 28). Global C-index: Harrell's concordance index computed over all segment pairs. Final-phase C-index: Harrell's C-index restricted to segment pairs both lying within the final 25% of the task, i.e. discrimination among segments close to exhaustion. Anticipation ($t_{80}$): time remaining before exercise cessation when the cumulative risk reached 80% of its final value, summarised as median [interquartile range] (Figure 3G). ≤2 min: participants whose warning fell within the final two minutes. Other values are mean ± SD.

| Segment | C-index | Anticipation $t_{80}$ (s) | PCs retained |
|---|---|---|---|
| 3 s | 0.891 ± 0.067 | 57 [32-73] | 15 ± 6 |
| 10 s | 0.897 ± 0.058 | 60 [33-73] | 6 ± 2 |
| 15 s | 0.903 ± 0.055 | 51 [31-67] | 5 ± 1 |
| 30 s | 0.910 ± 0.053 | 63 [36-82] | 4 ± 1 |
| 60 s | 0.926 ± 0.045 | 54 [33-91] | 3 ± 0 |

**Table S4. Cox proportional hazards model performance for EEG data (64 channels, 836 features) across segment lengths.** Values represent mean ± SD across participants (n = 28). C-index: Harrell's concordance index; Anticipation ($t_{80}$): time remaining before exercise cessation when the cumulative risk reached 80% of its final value, summarised as median [interquartile range] (Figure 3G). PCs: principal components retained per participant (70% variance threshold).

| **Segment** | **C-index** | **Anticipation $t_{80}$ (s)** |
|---|---|---|
| 3 s | 0.929 ± 0.022 | 38 [24-89] |
| 10 s | 0.933 ± 0.023 | 40 [23-94] |
| 15 s | 0.936 ± 0.024 | 38 [24-96] |
| 30 s | 0.944 ± 0.024 | 38 [28-100] |
| 60 s | 0.960 ± 0.024 | 44 [30-89] |

**Table S5. Cox proportional hazards model performance for cardiorespiratory data (5 features: $VO_2$, $VO_2$/kg, $VCO_2$, RER, HR) across segment lengths.** Values represent mean ± SD across participants (n = 28). C-index: Harrell's concordance index. Anticipation ($t_{80}$): time remaining before exercise cessation when the cumulative risk reached 80% of its final value, summarised as median [interquartile range]. Other values are mean ± SD.

| Segment | C-index | Anticipation $t_{80}$ (s) |
|---|---|---|
| 3 s | 0.945 ± 0.012 | 28 [20-50] |
| 10 s | 0.949 ± 0.014 | 27 [17-59] |
| 15 s | 0.952 ± 0.014 | 25 [18-57] |
| 30 s | 0.959 ± 0.017 | 32 [20-65] |
| 60 s | 0.972 ± 0.017 | 36 [26-54] |

**Table S6. Cox proportional hazards model performance for the combined model (early fusion: EEG + cardiorespiratory) across segment lengths. Values represent mean ± SD across participants (n = 28).** EEG features (64 channels, 836 features) were preprocessed with StandardScaler, winsorisation (±5 SD), and adaptive PCA (70% variance threshold); cardiorespiratory features ($VO_2$, $VO_2$/kg, $VCO_2$, RER, HR) were standardised and winsorised without PCA. Both feature sets were concatenated before fitting a single Ridge-penalised Cox model (penaliser = 0.1). C-index: Harrell's concordance index; Anticipation ($t_{80}$): time remaining before exercise cessation when the cumulative risk reached 80% of its final value, summarised as median [interquartile range]. Other values are mean ± SD.

| EEG window | PHY window | C-index | Anticipation $t_{80}$ (s) | ≤ 2min |
|---|---|---|---|---|
| 3 s | 30 s | 0.950 ± 0.015 | 25 [18-71] | 26/28 |
| 3 s | 60 s | 0.952 ± 0.015 | 31 [23-77] | 25/28 |
| 10 s | 30 s | 0.953 ± 0.016 | 28 [15-65] | 26/28 |
| 10 s | 60 s | 0.955 ± 0.016 | 32 [23-71] | 26/28 |
| 15 s | 30 s | 0.955 ± 0.016 | 27 [17-60] | 26/28 |
| 15 s | 60 s | 0.957 ± 0.016 | 34 [22-59] | 26/28 |
| 30 s | 60 s | 0.961 ± 0.018 | 34 [26-63] | 25/28 |

**Table S7. Cox proportional hazards model performance for the asynchronous combined model (EEG + cardiorespiratory, early fusion) across combinations of EEG and cardiorespiratory segment lengths.** EEG features were processed at finer temporal resolution (3, 10, 15, or 30 s) while cardiorespiratory features were extracted at coarser resolution (30 or 60 s); cardiorespiratory values were expanded to the EEG temporal grid by nearest-neighbour repetition before concatenation. Values represent mean ± SD across participants (n = 28). C-index: Harrell's concordance index; Anticipation ($t_{80}$): time remaining before exercise cessation when the cumulative risk reached 80% of its final value, summarised as median [interquartile range]. Other values are mean ± SD. ≤2 min: participants whose warning fell within the final two minutes. Other values are mean ± SD.

| Segment | ΔC-index | Stouffer Z (p) | Participants with p < 0.05 |
|---|---|---|---|
| **Whole exercise** | | | |
| 3 s | +0.016 | 17.6 ($9.0 \times 10^{-70}$) | 26/28 |
| 10 s | +0.016 | 17.1 ($3.5 \times 10^{-66}$) | 25/28 |
| 15 s | +0.017 | 16.6 ($2.6 \times 10^{-62}$) | 25/28 |
| 30 s | +0.015 | 12.3 ($8.2 \times 10^{-35}$) | 19/28 |
| 60 s | +0.012 | 6.0 ($7.8 \times 10^{-10}$) | 12/28 |
| **Final quarter of the exercise** | | | |
| 3 s | +0.080 | 14.0 ($6.6 \times 10^{-45}$) | 23/28 |
| 10 s | +0.099 | 11.7 ($9.7 \times 10^{-32}$) | 19/28 |
| 15 s | +0.111 | 10.5 ($3.9 \times 10^{-26}$) | 18/28 |
| 30 s | +0.090 | 7.0 ($1.6 \times 10^{-12}$) | 11/28 |
| 60 s | +0.116 | 1.3 ($9.5 \times 10^{-3}$) | 7/28 |

**Table S8. Permutation test of the neural contribution beyond cardiorespiratory monitoring across segment lengths.** For each participant the temporal order of the EEG block was permuted 10,000 times while the cardiorespiratory block was left intact, preserving the dimensionality and the marginal distribution of the neural features while destroying their alignment with the progression of the exercise. ΔC-index is the mean difference in concordance between the observed fused model and the permuted null. Stouffer Z combines the per-participant one-sided p values (n = 28). The final-quarter analysis restricts the concordance to pairs of segments both lying within the last 25% of the task. The neural contribution to the ranking of the whole exercise is significant at every segment length, and the magnitude of the gain in the final quarter is five to ten times that over the task as a whole. At the 60-s window the test remains significant, although with reduced power: only about six segments fall within the final quarter of the task.

| EEG window | Scheme | AUROC | AUPRC | F1 | n |
|---|---|---|---|---|---|
| 3 s | Random (stratified) folds | 0.990 ± 0.009 | 0.963 ± 0.030 | 0.918 ± 0.036 | 28 |
| 3 s | Blocked, gap 0 s | 0.956 ± 0.032 | 0.820 ± 0.136 | 0.821 ± 0.095 | 28 |
| 3 s | Blocked, gap 10 s | 0.950 ± 0.030 | 0.791 ± 0.140 | 0.811 ± 0.087 | 28 |
| 3 s | Blocked, gap 20 s | 0.940 ± 0.038 | 0.770 ± 0.154 | 0.780 ± 0.111 | 28 |
| 3 s | Blocked, gap 40 s | 0.926 ± 0.048 | 0.719 ± 0.170 | 0.749 ± 0.135 | 28 |
| 10 s | Random (stratified) folds | 0.981 ± 0.016 | 0.949 ± 0.034 | 0.922 ± 0.044 | 28 |
| 10 s | Blocked, gap 0 s | 0.942 ± 0.045 | 0.822 ± 0.126 | 0.831 ± 0.106 | 28 |
| 10 s | Blocked, gap 10 s | 0.949 ± 0.037 | 0.829 ± 0.129 | 0.836 ± 0.085 | 28 |
| 10 s | Blocked, gap 20 s | 0.932 ± 0.063 | 0.789 ± 0.148 | 0.805 ± 0.120 | 28 |
| 10 s | Blocked, gap 40 s | 0.911 ± 0.059 | 0.732 ± 0.145 | 0.767 ± 0.121 | 28 |
| 15 s | Random (stratified) folds | 0.979 ± 0.013 | 0.939 ± 0.036 | 0.918 ± 0.041 | 28 |
| 15 s | Blocked, gap 0 s | 0.932 ± 0.051 | 0.822 ± 0.124 | 0.839 ± 0.090 | 28 |
| 15 s | Blocked, gap 10 s | 0.924 ± 0.071 | 0.814 ± 0.143 | 0.835 ± 0.112 | 28 |
| 15 s | Blocked, gap 40 s | 0.895 ± 0.072 | 0.710 ± 0.177 | 0.775 ± 0.103 | 28 |
| 30 s | Random (stratified) folds | 0.951 ± 0.039 | 0.881 ± 0.085 | 0.891 ± 0.070 | 22 |
| 30 s | Blocked, gap 0 s | 0.906 ± 0.082 | 0.798 ± 0.141 | 0.824 ± 0.115 | 22 |
| 30 s | Blocked, gap 10 s | 0.890 ± 0.126 | 0.750 ± 0.178 | 0.787 ± 0.153 | 22 |

**Table S9. Binary classification of near- versus far-from-exhaustion EEG segments under random stratified folds and under blocked cross-validation with gaps of 0–40 s** (ten contiguous blocks per participant; training segments within the gap of the test block removed). Mean ± SD across participants (n = 28 at 3–15 s; n = 22 at 30 s). The 60 s window is omitted because too few segments remain per participant. Discrimination decreased only modestly as the gap widened (AUROC at 10 s: 0.981 random; 0.942, 0.949, 0.932 and 0.911 with gaps of 0, 10, 20 and 40 s). Gaps were converted to a whole number of segments (rounded, minimum one segment). At 15 s, gaps of 10 and 20 s both remove one segment (15 s), and at 30 s gaps of 10, 20 and 40 s all remove one segment (30 s); only distinct configurations are shown.

| EEG window | Scheme | Global C-index | Final-25% C-index | n |
|---|---|---|---|---|
| 3 s | In-sample (reference) | 0.891 ± 0.068 | 0.705 ± 0.108 | 28 |
| 3 s | Blocked, gap 0 s | 0.869 ± 0.088 | 0.638 ± 0.138 | 28 |
| 3 s | Blocked, gap 10 s | 0.867 ± 0.088 | 0.631 ± 0.142 | 28 |
| 3 s | Blocked, gap 20 s | 0.865 ± 0.087 | 0.623 ± 0.141 | 28 |
| 3 s | Blocked, gap 40 s | 0.860 ± 0.089 | 0.612 ± 0.151 | 28 |
| 10 s | In-sample (reference) | 0.897 ± 0.059 | 0.736 ± 0.107 | 28 |
| 10 s | Blocked, gap 0 s | 0.875 ± 0.067 | 0.678 ± 0.144 | 28 |
| 10 s | Blocked, gap 10 s | 0.871 ± 0.069 | 0.664 ± 0.157 | 28 |
| 10 s | Blocked, gap 20 s | 0.869 ± 0.069 | 0.656 ± 0.163 | 28 |
| 10 s | Blocked, gap 40 s | 0.865 ± 0.072 | 0.642 ± 0.175 | 28 |
| 15 s | In-sample (reference) | 0.903 ± 0.056 | 0.736 ± 0.112 | 28 |
| 15 s | Blocked, gap 0 s | 0.877 ± 0.069 | 0.683 ± 0.163 | 28 |
| 15 s | Blocked, gap 10 s | 0.874 ± 0.072 | 0.674 ± 0.165 | 28 |
| 15 s | Blocked, gap 40 s | 0.867 ± 0.076 | 0.649 ± 0.194 | 28 |
| 30 s | In-sample (reference) | 0.909 ± 0.055 | 0.743 ± 0.137 | 27 |
| 30 s | Blocked, gap 0 s | 0.880 ± 0.068 | 0.686 ± 0.187 | 27 |
| 30 s | Blocked, gap 10 s | 0.872 ± 0.072 | 0.659 ± 0.219 | 27 |

**Table S10. Within-participant EEG Cox concordance estimated out of sample**: models fitted on nine contiguous blocks and evaluated on the held-out block, with training segments within the indicated gap removed; C-index computed on pooled out-of-fold predictions. In-sample values are shown for reference. Mean ± SD (n = 28; 27 at 30 s). Gaps were converted to a whole number of segments (rounded, minimum one segment). At 15 s, gaps of 10 and 20 s both remove one segment (15 s), and at 30 s gaps of 10, 20 and 40 s all remove one segment (30 s); only distinct configurations are shown.

| Level | EEG | Cardiorespiratory | Combined | Friedman | Comb < Cardio | Comb < EEG | EEG vs Cardio | Dispersion p |
|---|---|---|---|---|---|---|---|---|
| 70% | 89 [54-102] | 59 [37-130] | 38 [27-78] | $\chi^2$=15.5, p=4.3 × $10^{-4}$ | 24/28, p < 0.001 | 21/28, p=0.009 | p=1.00 | 0.008 |
| 80% | 60 [33-73] | 40 [23-94] | 27 [17-59] | $\chi^2$=19.8, p=5.1 × $10^{-5}$ | 26/28, p < 0.001 | 21/28, p=0.012 | p=0.9 | 0.003 |
| 90% | 26 [16-33] | 27 [13-47] | 19 [10-36] | $\chi^2$=10.5, p=5.2 × $10^{-3}$ | 24/28, p < 0.001 | 17/28, p=0.24 | p=0.33 | 0.002 |

**Table S11. Anticipation (s, median [IQR]) when the cumulative risk reached 70%, 80% or 90% of its final value (10 s window, n = 28).** Friedman test across models; pairwise Wilcoxon signed-rank tests, Holm-corrected (number of participants in whom the combined warning came later); dispersion: sign-flip permutation test on absolute deviations from the median (10,000 permutations).

| Model | Within-participant C-index | LOPO C-index | LOPO C-index, median [range] | Participants above chance (C>0.5) |
|---|---|---|---|---|
| EEG | 0.897 ± 0.058 | 0.781 ± 0.081 | 0.799 [0.501-0.878] | 28/28 |
| Cardiorespiratory | 0.933 ± 0.023 | 0.916 ± 0.059 | 0.934 [0.688-0.966] | 28/28 |

**Table S12. Between-participant generalisation of the Cox survival model (10 s window, n = 28).** For each held-out participant, a ridge-penalised Cox model was trained on the remaining 27 participants (leave-one-participant-out, LOPO), with standardisation, winsorisation and, for EEG, the 70%-variance PCA fitted on the training participants only, and evaluated on the held-out participant. Within-participant values (models fitted on each participant's own data; Table S3) are shown for reference. C-index: Harrell's concordance index. Mean ± SD and median [range] across held-out participants. The cardiorespiratory model transferred almost intact across individuals, whereas EEG concordance fell markedly, indicating that the cortical signature of approaching exhaustion is largely participant-specific (see Fig. S17B and C).

| | Planned | Implemented |
|---|---|---|
| ***Hypotheses and aims*** | | |
| Primary hypothesis (H1) | Lempel–Ziv complexity and wSMI drop abruptly once the anaerobic threshold is exceeded, with an abrupt and diffuse loss of connectivity. | Complexity, including Lempel–Ziv complexity, decreased as physiological strain rose. wSMI did not decrease overall. It increased at the two longer lags and decreased at the shortest one. The anaerobic threshold was not estimated. The analyses went directly to the point of exhaustion, with the neural measures modelled against a continuous index of physiological strain, so no abrupt change at the threshold was tested. |
| H2 | Reduced integration (graph theory) and complexity at the ventilatory threshold. | The ventilatory threshold was not estimated. As for H1, the analyses went directly to the point of exhaustion rather than to the physiological thresholds. |
| H3–H5 | Multivariate decoding of the auditory task (tone, match and mismatch, omissions) and its generalisation across effort levels. | Not reported here. The auditory task addresses different questions and is reported separately. |
| Prediction of exhaustion | Stated as an aim, without a specified method. | Addressed with participant-specific classifiers and Cox models, which were not specified in the preregistration. |
| Phenomenological questionnaire | Planned. | Not administered, to avoid lengthening the session for participants. |
| ***Methods*** | | |
| Sample | 30 expert cyclists aged 18–36 years. | 32 recruited and 28 analysed (4 women), aged 18–52 years. |
| Starting load | 120 W, increasing by 30 W every 5 min. | 120 W for men and 100 W for women, increasing by 30 W every 5 min. |
| End of the test | Cadence below 45 rpm. | Power output more than 10 SD below its mean over the final 2 min, identified offline. |
| Resting periods | Eyes-open rest without tones, then with tones. | Eyes-open rest with tones before and after exercise. Eyes-closed rest was also recorded at the beginning and end of the session but not analysed. |

| | | |
|---|---|---|
| Epoch rejection | Epochs exceeding ±50 dB (0–2 Hz) or ±100 dB (20–40 Hz) rejected. | Thresholds of +25 and −100 dB in the 20–40 Hz band, and flagged epochs replaced by the preceding clean epoch to keep the recording continuous. |
| Muscle artefacts | EEGLAB and MoBILAB routines. | No MoBILAB. Band-pass filter between 1 and 40 Hz. |
| Neural measures | Lempel–Ziv complexity and wSMI as primary measures, possibly with Kolmogorov complexity, wPLI, detrended fluctuation analysis, sample entropy, graph theory and multivariate decoding. | Oscillatory power and the aperiodic component (not preregistered), seven complexity measures and wSMI at four lags. No wPLI, detrended fluctuation analysis or decoding. |
| Statistics | Per-participant effect sizes, described rather than tested. | Piecewise linear models with permutation-based family-wise correction, gradient-boosted classifiers, Cox models, leave-one-participant-out validation and permutation null models. |

**Table S13. Deviations from the preregistered protocol.** Comparison between the preregistration (C. Avancini, C. Alameda, D. Sanabria, T. Bekinschtein, S. van Gaal, L. F. Ciria, Characterising the neural dynamics of physical exhaustion, Open Science Framework (2024); https://doi.org/10.17605/OSF.IO/TW5KV), registered before data collection, and the study reported here.
